# GALE-dependent glycoproteome remodelling is a determinant of oncogenic RAS transformation

**DOI:** 10.64898/2026.06.15.732364

**Authors:** Frauke Stölting, Rita Vesce, César Rodríguez-Santana, Marina Ciscar, Alberto Delaidelli, Qiaochu Lin, Christian Ballmeyer, Sophie Chan, Ramona Lesch, Ruitong Li, Abhimanyu DasGupta, Bin Yu, Daniel Picard, Vithusan Suppiyar, Francesco Casini, Aslihan Yavas, Sami A. Safi, Wolfram Trudo Knoefel, Lena Häberle, Marc Remke, David Koppstein, W. Brent Derry, Poul H. Sorensen, Guido Reifenberger, Gabriel Leprivier, Georg Fluegen, Irene Esposito, Naiara Santana-Codina, Marc D. Driessen, Ulla I.M. Gerling-Driessen, Jonathan K. M. Lim

## Abstract

Oncogenic transformation is accompanied by extensive remodelling of the cellular glycosylation landscape, yet the mechanisms linking oncogenic signalling to glycoproteome reprogramming remain poorly defined. We used site-resolved intact glycoproteomics to systematically map oncogenic RAS-dependent changes across the N- and O-linked glycoproteome. Integrating multiomics profiling with functional approaches, we identify UDP-glucose 4-epimerase (GALE) as a transcriptional target of oncogenic RAS that is induced through MAPK signalling and MYC-dependent transcription. GALE depletion selectively disrupted RAS-dependent glycoproteome remodelling, preferentially impairing sialylation of N-linked glycans and O-linked GalNAc-type glycosylation. Functionally, these glycosylation defects were accompanied by suppressed anchorage-independent growth and reduced in vivo tumorigenicity of *KRAS*-mutant cells, establishing GALE-dependent nucleotide-sugar interconversion as a requirement for malignant growth. Clinically, GALE expression was elevated in pancreatic ductal adenocarcinoma, wherein *KRAS* mutations are highly prevalent, and across a broad spectrum of human cancers. Collectively, our findings define a regulatory axis that connects oncogenic signalling to nucleotide-sugar precursor availability and diversification of the cellular glycan repertoire, revealing GALE as a metabolic dependency in RAS-driven cancer.

## Introduction

Oncogenic activation of the RAS family of small GTPases represents one of the most pervasive drivers of human malignancy, with *KRAS*, *HRAS*, and *NRAS* mutations collectively occurring in over 25% of all cancers^1^. Among these, *KRAS* mutations are most frequent and found in multiple tumor entities including pancreatic cancer, colorectal cancer, and non-small cell lung cancer (NSCLC). In physiological settings, RAS proteins relay extracellular growth signals to intracellular pathways controlling proliferation, differentiation, and survival. Oncogenic mutations lock RAS in a constitutively active state, thereby driving sustained mitogenic signalling and tumor initiation. For decades, RAS proteins were considered as “undruggable,” due to its lack of suitable binding pockets, while attempts to mislocalise RAS or target its effectors were limited by toxicity or lack of efficacy^2^. Recent successes with covalent inhibitors targeting specific KRAS variants have demonstrated that RAS can be therapeutically targeted; however, clinical responses in patients remain transient and are eventually followed by drug resistance^3,4^. These limitations underscore the need to better define the downstream dependencies and metabolic rewiring induced by oncogenic RAS, both to understand tumor biology and to identify novel therapeutic vulnerabilities.

Glycosylation is one of the most abundant and structurally diverse post-translational modifications of proteins, regulating a wide range of biological processes including protein folding, trafficking, receptor signalling, and cell–cell interactions^5^. Glycosylation is orchestrated by a complex network of glycosyltransferases, glycosidases, nucleotide sugar transporters, and metabolic pathways that generate activated sugar donors. Aberrant glycosylation is a common feature of malignant progression, with tumors exhibiting characteristic changes including increased branched and fucosylated N-glycans, truncated O-linked glycans, increased sialylation, and the emergence of tumor-associated carbohydrate antigens (TACAs) such as sialyl-Lewis A^6,7^.

A longstanding observation is that oncogenic transformation, including that driven by mutant RAS, is concomitant with widespread remodelling of the cellular glycosylation landscape^8–10^. A seminal study by Ying and colleagues demonstrated that oncogenic KRAS enhances glucose flux into the hexosamine biosynthesis pathway (HBP), thereby increasing the production of UDP-N-acetylglucosamine (UDP-GlcNAc), a central substrate for glycosylation^11^. Inhibition of the rate-limiting enzyme of HBP, Gfpt1, led to the impairment of tumor growth in KRAS-mutant models, suggesting the critical importance of glycosylation for tumor progression. While this early work established a direct mechanistic link between oncogenic signalling and the metabolic supply of glycosylation precursors, how oncogenes coordinate the diversification and specificity of tumor-associated glycan composition and topographies remains unresolved. Understanding the complex interplay between oncogenic drivers and global changes to glycosylation could provide new avenues for RAS-specific diagnostic and therapeutic development.

We employed combinatorial approaches to interrogate how oncogenic RAS reprograms the glycoproteome and uncovered a previously unrecognized role of the UDP-glucose 4-epimerase (GALE) as a central effector of this process. We show that RAS-MAPK signalling, through MYC activity, transcriptionally upregulates *GALE*, thereby selectively enhancing galactose- and GalNAc-dependent glycosylation, including sialylation. Using intact glycoproteomics, we further present systematic site-specific precision mapping of the GALE-dependent glycoproteome in engineered and established KRAS-mutant cell models. Functional assays demonstrate that GALE-dependent glycosylation is a cell-intrinsic determinant of oncogenic transformation and tumor growth in vivo. Moreover, GALE expression is elevated in *KRAS*-mutant cancers and correlates with disease severity across multiple tumor types, suggesting a broader clinical relevance. Our findings define a mechanistic link between oncogenic RAS signalling and selective glycosylation remodelling and point to GALE as a potential therapeutic vulnerability.

## Results

### Oncogenic RAS transformation leads to selective remodelling of the glycoproteome

To understand the impact of oncogenic transformation on the cellular glycosylation landscape, we used NIH 3T3 murine fibroblasts stably transformed with *KRAS^G12V^*(henceforth referred to as NIH 3T3 KRAS^G12V^), which we previously leveraged to identify novel downstream effectors of oncogenic RAS transformation^12,13^. We subjected oncogenic KRAS-transformed and non-transformed cells to unbiased, site-specific, N- and O-linked glycoproteomic profiling using the strong anion exchange, electrostatic repulsion and hydrophilic interaction chromatography (SAX-ERLIC) enrichment strategy, with subsequent LC-MS/MS analysis^14^. Using this approach, we catalogued 2,766 unique glycoforms (GlycoIDs) in NIH 3T3 KRAS^G12V^ cells, corresponding to 538 glycoproteins containing 897 unambiguous glycosites detected from a total of 12,138 glycopeptide-spectrum matches (glycoPSM) (Fig. S1A, Table S1). In non-transformed NIH 3T3 cells, i.e. NIH 3T3 expressing murine stem cell virus (MSCV) control vector (henceforth referred to as NIH 3T3 MSCV), 2,875 unique GlycoIDs corresponding to 539 glycoproteins containing 946 glycosites were detected from 12,014 glycoPSMs (Fig. S1B and Table S1). From these data, we identified 503 N-linked glycoproteins that displayed a shift in glycan abundances between KRAS^G12V^-transformed cells relative to non-transformed counterparts (Fig. 1A and Table S1). Overall, sialic acid-containing N-glycans [including sialyl-fucosylated (Sia-Fuc) structures] were the most enriched glycan class in KRAS^G12V^-transformed cells. By contrast, hybrid-type and high-mannose showed the greatest net depletion, although high-mannose-modified N-linked glycoproteins exhibited bidirectional changes (Fig. 1A and B). Using lectin profiling as an orthogonal approach, we found that *Sambucus nigra* II (SNA), specific for α2,6 sialylated structures, showed higher binding in KRAS^G12V^-transformed versus non-transformed cells (Fig. 1C). Amongst the glycoproteins showing the most changes in sialic acid glycans were those involved in lysosome/endosome trafficking (IGF2R, LAMP1 and 2, TPP1, CTSD, SCARB2, M6PR, SYPL1, CPD, MAN2A1, GAA), extracellular matrix organization (FBLN2, BGN, HS2ST1, FN1, THBS1), integrin/cell-adhesion signalling (VCAM1, NPTN, ALCAM, EPHA2, ITGA2, ITGA6, ADGRA2, ITGB1, PTK7), and scavenger/endocytic receptors (LRP1, MRC2, SCARA2, COLEC12) (Fig. 1D). Interestingly, many of these glycoproteins showed significant heterogeneity in glycan changes upon KRAS^G12V^-mediated transformation (Fig. S1, C and D). For example, low-density lipoprotein receptor-related protein 1 (LRP11), integrin β1 (ITGB1) and fibronectin 1 (FN1) were among the glycoproteins most enriched for sialylated glycans, yet showed a concomitant loss of high-mannose structures (Table S1). To gain a more comprehensive understanding of the cellular processes altered in response to oncogenic KRAS-mediated glycoproteome remodelling, we performed pathway enrichment and protein-protein interaction clustering analysis on all differentially regulated N-linked glycoproteins^15^. Enriched pathways associated with oncogenic KRAS transformation included extracellular matrix organization, cell-cell adhesion, wound healing, and blood vessel organization, while protein-protein interaction (PPI) networks included integrin signalling, ER stress response, collagen biosynthesis, semaphorin-plexin signalling, lysosome/endosome trafficking, ECM remodelling, and N-glycan biosynthesis (Fig. 1E and Fig. S1E).

**Fig. 1.**
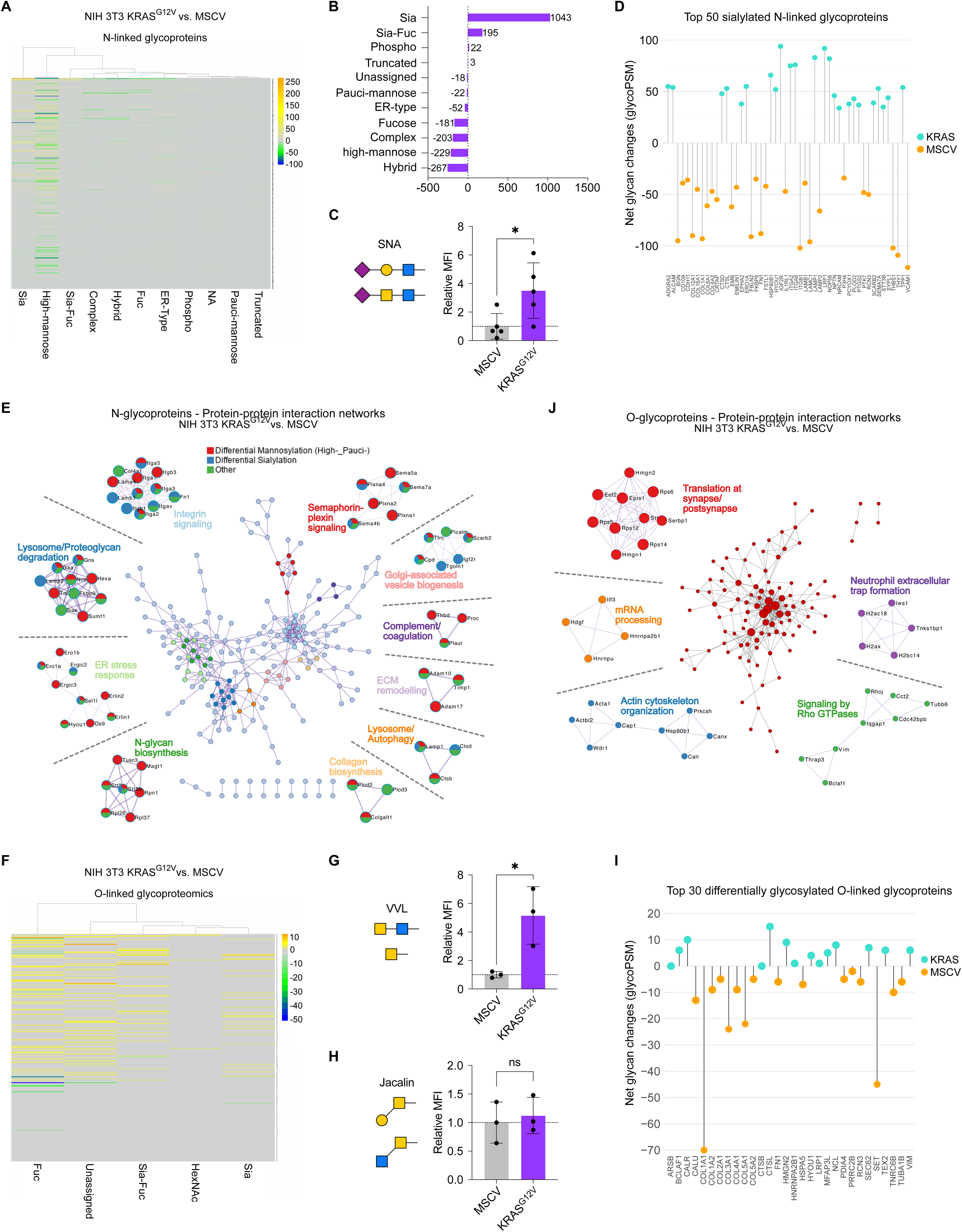
Oncogenic RAS transformation leads to selective remodelling of the glycoproteomic landscape. (A) Differential glycan analysis of N-linked glycoproteins comparing NIH 3T3 cells expressing mutant KRAS^G12V^ (NIH 3T3 KRAS^G12V^) versus control MSCV, obtained via SAX-ERLIC enrichment method and tandem mass spectrometry. Counts represent glycopeptide-spectrum matches (glycoPSM). Sia represents sialylation; Sia-Fuc represents Sialyl-fucosylated; NA represents not allocated to any of the indicated glycan types. (B) Sum total glycoPSM of each glycan type across all N-linked glycoproteins, comparing NIH 3T3 KRAS^G12V^ versus control MSCV. (C) SNA-fluorescein labelling of NIH 3T3 KRAS^G12V^ and MSCV cells. Relative MFI (mean fluorescence intensity) = [MFI_with lectin_-MFI_without lectin_]/MFI_without lectin_ normalized to the control from each biological replicate. (D) Net glycan changes (glycoPSM) across all glycan types for each of top 50 positively-sialylated N-linked glycoproteins comparing NIH 3T3 KRAS^G12V^ to MSCV. (E) Protein-protein interaction (PPI) analysis of N-linked glycoproteins with positive glycosylation changes in NIH 3T3 KRAS^G12V^ versus MSCV cells, with the contribution of high- and pauci-mannosylated N-glycoproteins, sialylated (including Sia-Fuc), and other glycan type indicated. The center network represents the full interactome, and the surrounding networks indicated representative functions. (F) Differential glycan analysis of O-linked glycoproteins comparing NIH 3T3 KRAS^G12V^ versus MSCV, obtained as in (A). Fuc represents fucosylated; HexNAc represents single O-linked HexNAc; Unassigned represents general O-linked glycan. (G) VVL-fluorescein and (H) Jacalin-fluorescein labeling of NIH 3T3 KRAS^G12V^ and MSCV cells, with relative MFI calculated as in (C). (I) Net glycan changes across all glycan types (glycoPSM) for O-linked glycoproteins showing the most significant glycosylation changes (based on total absolute glycoPSM) comparing NIH 3T3 KRAS^G12V^ to MSCV. (J) PPI analysis of O-linked glycoproteins with positive glycosylation changes in NIH 3T3 KRAS^G12V^ versus MSCV cells. The center network represents the full interactome, and the surrounding networks indicated representative functions. Unless otherwise indicated, data represent mean ± SD of at least three independent experiments. Unpaired, two-tailed *t test*; \**P* < 0.05, \*\**P* < 0.01, \*\*\**P* < 0.001, \*\*\*\**P* < 0.0001. ns, not significant.

We additionally profiled 273 O-linked glycoproteins exhibiting glycan changes in oncogenic KRAS-transformed cells (Table S2). Overall, we observed a general enrichment across all identified O-linked glycan classes in KRAS^G12V^-transformed cells, consistent with increased binding of *Vicia villosa* (VVL) lectin, specific for Tn, core 5 and core 7 O-glycan structures (Fig. 1F and G, Fig. S1F)^16^. Interestingly, we did not observe any changes in the binding of *Jacalin* lectin, which is specific for core 1 and core 3 O-linked glycans, and sialyl-T antigen (Fig. 1H). However, this may be due to the appending of additional inhibitory glycan moieties onto terminal glycans otherwise recognized by *Jacalin*. We also observed a concomitant decrease in *Wheat germ agglutinin* (WGA) lectin binding, suggesting that the enrichment of O-linked glycans observed are more likely to be O-GalNAc glycosylation rather than the post-translational modification – O-GlcNAcylation (Fig. S1G). Enriched pathways or PPI networks representing differentially regulated O-linked glycoproteins included collagen formation, cell-cell communication, Rho GTPase signalling (BCLAF1, VIM), actin cytoskeleton organization (CALR), mRNA processing (HNRNPA2B1), and translation at synapse (HMGN2), among others (Fig. 1I and J, Fig. S1H). Many of these pathways and interaction networks, as well as the individual glycoproteins that constitute these networks, have been well-described to contribute to the tumorigenic process, supporting the idea that O-linked glycosylation might constitute an additional important layer of regulation of oncogenic phenotypes. Taken together, our glycoproteomic survey demonstrates a level of regulatory specificity in glycan composition that occurs following oncogenic KRAS transformation, suggesting selective remodelling of the glycoproteome.

### Oncogenic RAS signalling drives GALE upregulation

To identify novel mediators regulated by oncogenic KRAS signalling that directly contribute to a rewired glycan landscape, we leveraged our previously reported integrated proteo-transcriptomic approach performed on KRAS^G12V^-transformed versus non-transformed cells^13,17^. Interestingly, none of the candidate genes/proteins upregulated by mutant KRAS had annotated function in glycosylation or glycosylation-associated metabolic pathways, except *GALE*, which encodes UDP-glucose 4’-epimerase (Fig. 2A). Orthogonal approaches confirmed the upregulation of GALE expression in NIH 3T3 KRAS^G12V^ cells relative to isogenic control cells, both at the transcript and protein levels (Fig. 2B). GALE catalyzes the epimerization of two pairs of nucleotide sugars, UDP-glucose and UDP-galactose, and in an orthogonal reaction their corresponding N-acetyl-hexosamines, UDP-GlcNAc and UDP-GalNAc, ensuring supply of these donor substrates for glycosylation reactions (Fig. 2C). Thus, we hypothesized that GALE might facilitate oncogenic RAS-mediated remodelling of the glycan landscape. Human BJ fibroblasts serially immortalized with the catalytic subunit of human telomerase (hTERT) and SV40 large T antigen (LT) also showed an increase in GALE expression following ectopic expression of oncogenic *HRAS^G12V^*, suggesting that oncogenic RAS-mediated regulation of GALE is not isoform-specific or mutant-specific (Fig. 2D). Similarly, human mammary epithelial cells (HMEC) expressing *HRAS^G12V^*showed the highest induction of *GALE* expression relative to control, or relative to other oncogenes including *CTNNB1* (encoding β-catenin), and *SRC,* suggesting that the regulation of *GALE* by oncogenic signalling is specific to RAS (Fig. 2E). We next turned to published microarray data from a doxycycline (dox)-inducible *KRAS^G12D^* pancreatic ductal adenocarcinoma (PDAC) mouse model previously reported by Ying and colleagues, termed the iKras model^11^. Consistently, dox withdrawal, corresponding to *KRAS^G12D^* extinction, led to a decrease in *GALE* expression in vitro and in vivo (Fig. 2, F and G). To understand the cellular distribution of *GALE* expression by oncogenic RAS in tumors, we next queried single-cell RNA sequencing data from a similar *Ptf1a-CreER*, *LSL-KRAS^G12D^, LSL-tdTomato* mouse model of PDAC reported by Schlesinger et al^18^. This analysis revealed that *GALE* expression was largely confined to malignant tumor cells, showing significant overlap with *tdTomato* signal representing KRAS^G12D^ transgene expression (Fig. 2H). To extend our findings into clinically relevant cell models, we used shRNA to ablate *KRAS* expression in two KRAS-mutant human cancer cell lines, namely the non-small cell lung carcinoma (NSCLC) cell line NCI-H460 and the PDAC cell line CFPAC1, which in both instances led to decreases in *GALE* expression (Fig. 2I and Fig. S2, A and B). In agreement, pharmacological inhibition of KRAS^G12C^ in a *KRAS^G12C^*-expressing PDAC cell line, MIA PaCa-2, with either AMG-510 or ARS-1620 effectively decreased *GALE* expression (Fig. 2J and Fig. S2C). To assess the generalizability of oncogenic RAS in regulating *GALE* expression, we performed a pan-cancer analysis of human cancer cell lines from the Cancer Cell Line Encyclopedia (CCLE) dataset, which revealed that cells harboring *KRAS* mutations (*KRAS^mut^*) were associated with higher *GALE* mRNA expression, as compared to *KRAS* wild-type (*KRAS^wt^*) cell lines (Fig. 2K). Since RAS pathway activation could occur in cancer cells even in the absence of *KRAS* mutations, we further analyzed RAS pathway activity in the same dataset using a validated transcriptional signature of 84 genes (RAS84) optimized to capture RAS oncogenic activity^19^. Consistent with our previous data, RAS pathway activation was also significantly correlated with *GALE* mRNA expression (Fig. 2L). Together, our data suggest that oncogenic RAS signalling leads to transcriptional upregulation of *GALE*.

**Fig. 2.**
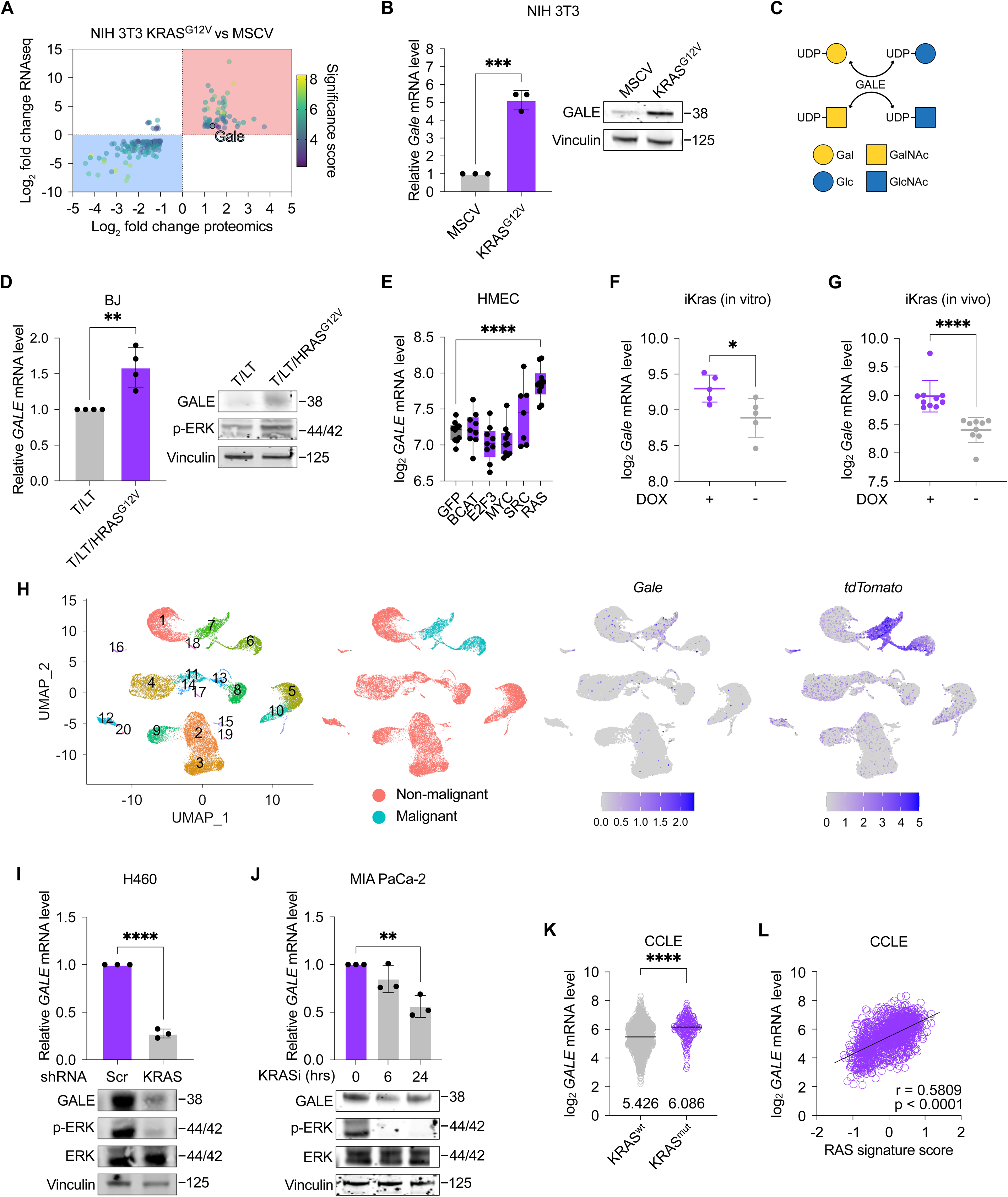
Oncogenic RAS transformation leads to transcriptional upregulation of GALE (A) Volcano plot summarizing fold changes of differentially regulated genes/proteins from RNA sequencing or proteomics of NIH 3T3 cells expressing mutant KRAS^G12V^ versus control MSCV. Significance score is derived from the sum of -log_10_ individual p-values from each platform. RNA sequencing and proteomics data were reanalyzed from previous studies^13,17^. (B) Relative *Gale* mRNA expression analyzed by qRT-PCR, and GALE protein expression analyzed by Western blot in NIH 3T3 cells expressing mutant KRAS^G12V^ or control MSCV. (C) Schematic depicting the molecular function of GALE. (D) Relative *GALE* mRNA expression analyzed by qRT-PCR, and protein levels of GALE and phosphorylated ERK (p-ERK) analyzed by Western blot in BJ cells expressing human telomerase (hTERT), SV40 large T (LT) and mutant *HRAS^G12V^* (hTERT/LT/HRAS^G12V^) or control hTERT/LT. (E) Relative *GALE* mRNA expression in human mammary epithelial cells (HMEC) expressing indicated oncogenes or GFP control, analyzed by microarray. Data are obtained from GSE3151, accessed via the R2 platform (http://r2.amc.nl)^21^. Relative *Gale* mRNA expression in doxycycline-inducible *KRAS* (iKRAS) model of PDAC (F) in vitro, and (G) in vivo. ON represents doxycycline-treated iKRAS cells or mice, while OFF represents doxycycline withdrawal from iKRAS cells or mice, obtained from GSE32277 via NCBI-GEO platform^11^. (H) UMAP of single cell RNA sequencing data from pancreatic tissue of a murine PDAC model (Ptf1a^CreER^;LSL-Kras^G12D^;LSL-*tdTomato*) showing malignant cell clusters, and expression of *GALE* and *tdTomato*^18^. (I) Relative *GALE* mRNA expression analyzed by qRT-PCR and GALE, p-ERK, and ERK protein levels analyzed by Western blot in shKRAS knockdown and scrambled (scr) control H460 cells. (J) Relative *GALE* mRNA expression analyzed by qRT-PCR and GALE, p-ERK, and ERK protein levels analyzed by Western Blot, in MIA PaCa-2 cells upon treatment with AMG510 (10 µM) or vehicle for the indicated times. (K) Correlation between KRAS mutation status and GALE expression in cancer cells. *wt, wild-type. mut, mutant.* The numbers indicate the median for each column. (L) Correlation between RAS pathway activation signature score and GALE expression in cancer cells. Correlation was quantified using Spearman’s rank correlation coefficient. Data are obtained from the Cancer Cell Line Encyclopedia (CCLE) database, accessed via the DepMap portal^19,51^. Unless otherwise indicated, data represent mean ± SD of at least three independent experiments. Unpaired, two-tailed *t test*; \**P* < 0.05, \*\**P* < 0.01, \*\*\**P* < 0.001, \*\*\*\**P* < 0.0001. ns, not significant.

### RAS/MAPK and MYC mediate *GALE* transcriptional upregulation downstream of mutant RAS

We next sought to determine the signalling and transcriptional mechanisms by which oncogenic RAS signalling regulates *GALE* expression. We focused on two canonical signalling axes downstream of RAS, namely RAF-MEK-ERK and PI3K-AKT, and found that treatment with the MEK inhibitor PD184352 but not the AKT inhibitor MK-2206 led to decreased Gale expression, concurrent with the suppression of phosphorylated ERK (p-ERK), suggesting that transcriptional regulation of *GALE* downstream of oncogenic RAS is mediated by MAPK (RAF-MEK-ERK) signalling (Fig. 3A and Fig. S3A). Similar effects were observed with MEK inhibition in *KRAS*-mutant cells derived from murine lung adenocarcinoma tumors of conditional Cas9-C57BL/6 mice intubated with adeno-associated viruses (AAV) to induce *KRAS* transgene expression, henceforth referred to as KP cells, as well as a *KRAS^G12D^*-mutant human PDAC cell line, PANC-1 cells (Fig. 3B and Fig. S3B)^20^.

**Fig. 3.**
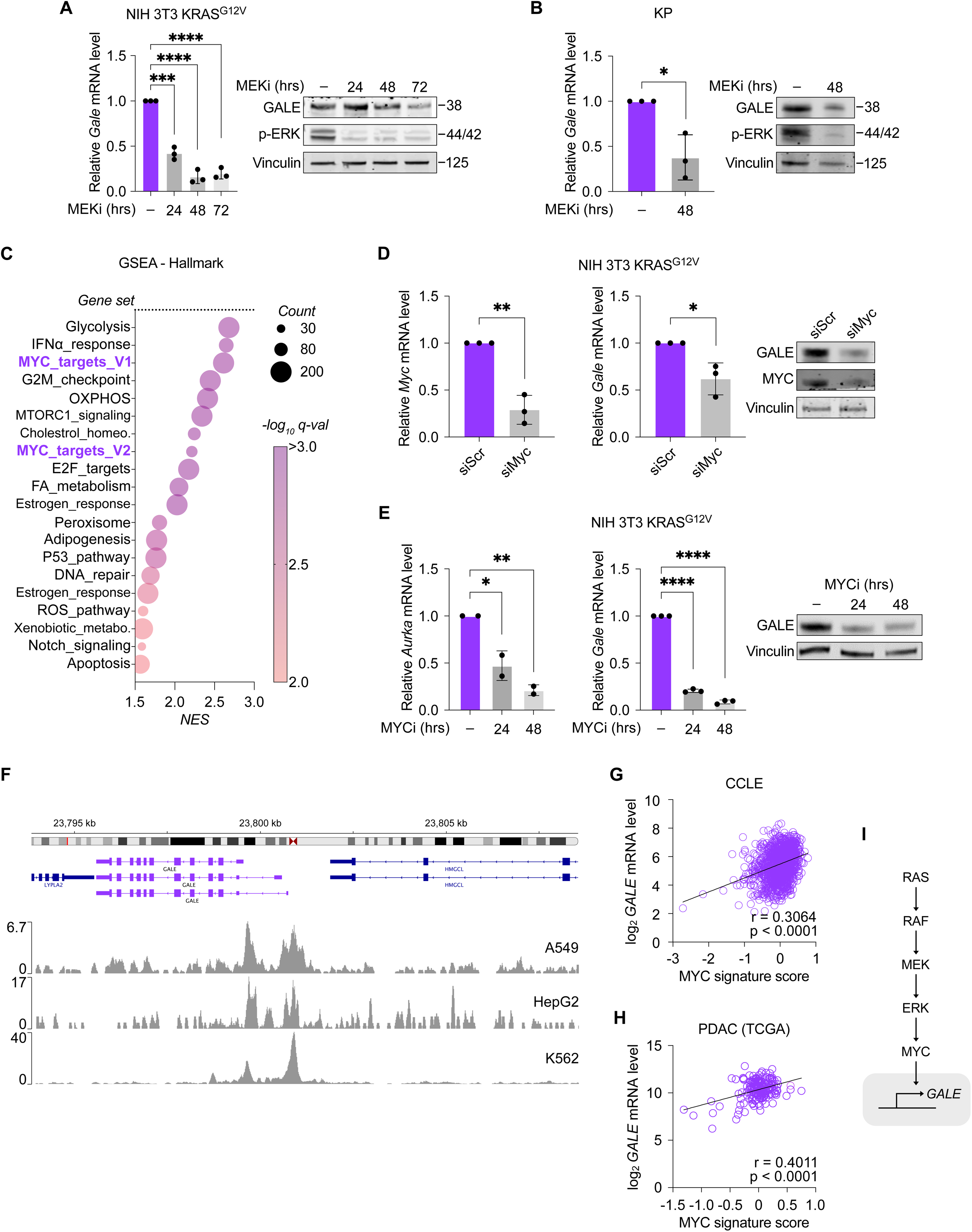
GALE induction downstream of oncogenic RAS is regulated by MAPK signalling and the transcription factor MYC. Relative *Gale* mRNA expression analyzed by qRT-PCR, and protein expression of GALE and p-ERK analyzed by Western Blot in (A) NIH 3T3 cells expressing mutant KRAS^G12V^ (NIH 3T3 KRAS^G12V^) and (B) KP cells upon treatment with MEK inhibitor PD184352 (MEKi; 10 µM) or vehicle for the indicated time. (C) Gene set enrichment analysis (GSEA) for Human Molecular Signatures Database (MSigDB) Hallmark gene sets performed on genes positively co-expressed with GALE in pancreatic ductal adenocarcinoma (PDAC) from The Cancer Genome Atlas (TCGA), analyzed using the R2 platform. (D) Relative *Gale* and *Myc* mRNA expression analyzed by qRT-PCR, and protein expression of MYC and GALE analyzed by Western Blot in NIH 3T3 KRAS^G12V^ cells transfected with siRNA against *Myc* (siMyc) or scrambled (siScr) control. (E) Relative *Gale* and *Aurka* mRNA expression analyzed by qRT-PCR, and protein expression of GALE analyzed by Western Blot in NIH 3T3 KRAS^G12V^ cells treated with MYC/MAX inhibitor 10058-F4 (MYCi; 10 µM) or vehicle for the indicated times. (F) ChIP-sequencing data showing MYC occupancy profiles at the GALE locus in the indicated cell lines. Data are obtained from UCSC Encode database, analyzed using Integrative Genomics Viewer (IGV)^52^. Correlation between GALE mRNA expression and MYC signature core in (G) cancer cells from the CCLE database, accessed via the DepMap portal and (H) PDAC samples from TCGA^21^. Correlation was quantified using Spearman’s rank correlation coefficient. (I) Schematic depicting the suggested mechanistic model for GALE regulation downstream of oncogenic RAS signalling. Unless otherwise indicated, data represent mean ± SD of at least three independent experiments. Unpaired, two-tailed *t test*; \**P* < 0.05, \*\**P* < 0.01, \*\*\**P* < 0.001, \*\*\*\**P* < 0.0001. ns, not significant.

To identify lead transcription factors that drive *GALE* transcription downstream of oncogenic RAS signalling, we rank-sorted PDAC samples from TCGA into high and low *GALE* tumors and performed gene set enrichment analysis (GSEA) on differentially-expressed genes from the two groups. This analysis revealed that c-MYC (MYC) target genes are among the most significantly enriched in *GALE*-high versus *GALE*-low samples (Fig. 3C). Supervised GSEA on *GALE*-high versus *GALE*-low PDAC tumors, or *GALE*-high versus *GALE*-low cell lines from CCLE similarly demonstrated that multiple orthogonal gene sets corresponding to MYC targets showed significant enrichment (Fig. S3C and D). Given this initial evidence, we decided to pursue MYC as the transcription factor candidate mediating *GALE* expression downstream of oncogenic RAS signalling, which to our knowledge has not been reported. We experimentally assessed the regulation of GALE expression by MYC and showed that siRNA-mediated knockdown of *Myc* in NIH 3T3 KRAS^G12V^ significantly decreased GALE expression (Fig. 3D). Similarly, knockdown of *Myc* in KP, or of *MYC* in the *KRAS*-mutant PDAC line PaTu-8988T led to a significant decrease in *GALE* expression, albeit only at the transcriptional level, possibly due to the stability of GALE protein in these lines (Fig. S3, E and F). Pharmacological inhibition of MYX/MAX dimerization using 10058-F4 also led to reduced GALE expression in NIH 3T3 KRAS^G12V^, along with reduced levels of *Aurka*, a well-characterized MYC target gene (Fig. 3E). Further, chromatin immunoprecipitation (ChIP)-sequencing analyses from publicly-available UCSC Encode data showed that endogenous MYC appeared to be among the transcription factors occupying transcriptionally active regions of the *GALE* promoter in three distinct cancer cell lines (Fig. 3F). Finally, *GALE* expression was significantly associated with a MYC signature both in cell lines from CCLE as well as clinical samples from multiple PDAC cohorts (Fig. 3, G and H; Fig. S3, G and H)^21^. Together, our data suggest that RAS/MAPK signalling and MYC function mediate oncogenic RAS regulation of GALE expression (Fig. 3I).

### GALE contributes to glycoproteome remodelling in oncogenic RAS-transformed cells

To ascertain whether GALE functionally contributes to remodelling of the glycoproteome in oncogenic RAS-transformed cells, we first generated *Gale*/*GALE* knockdown cell models in NIH 3T3 KRAS^G12V^ and PANC-1 cells using CRISPR-interference (CRISPRi). Namely, two single guide RNA (sgRNA) constructs targeting *Gale* (sgGale-1 and sgGale-2) or *GALE* (sgGALE-1 and sgGALE-2) were compared to a control sgRNA targeting EGFP (sgEGFP) (Fig. S4A and B). We performed parallel SAX-ERLIC-enrichment and tandem MS-based N- and O-linked glycoproteomic profiling on GALE-depleted lines from both cell models. From this, we catalogued a total of 2,158 unique glycoforms (GlycoIDs) in NIH 3T3 KRAS^G12V^ cells (GALE-depleted and control) corresponding to 548 glycoproteins containing 895 unambiguous glycosites detected from a total of 5,335 glycopeptide-spectrum matches (glycoPSM) (Fig. S4C and D, Table S3) and a total of 4,629 unique glycoforms (GlycoIDs) in PANC-1 (GALE-depleted and control) cells, corresponding to 1,231 glycoproteins containing 2,139 unambiguous glycosites detected from a total of 18,185 glycopeptide-spectrum matches (glycoPSM) (Fig. S4E and F, Table S4). As UDP-glucose and UDP-GlcNAc originate from glucose metabolism, we hypothesize that GALE inhibition should limit their conversion into UDP-galactose and UDP-GalNAc, thereby disrupting galactose or GalNAc-dependent glycosylation, as well as sialylation, since sialic acids are predominantly added to terminal galactose, GalNAc, or sialic acid residues. In support of our hypothesis, many N-linked glycoproteins showed a decrease in sialylation in *GALE* knockdown PANC-1 cells relative to controls, and in *Gale* knockdown NIH 3T3 KRAS^G12V^ relative to controls, consistent with SNA lectin binding profiles, suggesting that loss of GALE activity indirectly limits sialylation by reducing these underlying acceptor substrates (Fig. 4, A-D, Fig. S4, G and H). These findings are in agreement with our initial observation that oncogenic RAS transformation increases N-linked sialylation, suggesting that GALE mediates a substantial fraction of these changes. In agreement, of the 503 N-linked glycoproteins altered upon oncogenic KRAS transformation in NIH 3T3 cells, 218 (43.3%) and 211 (41.9%) also showed glycosylation changes following *Gale* knockdown in NIH 3T3 KRAS^G12V^ cells and *GALE* knockdown in PANC-1 cells respectively (Fig. S4, I and J). Beyond this overlap at the level of individual glycoproteins, there was significant concordance between their associated GO terms, suggesting that glycosylation-related pathways remodelled during oncogenic KRAS transformation are recurrently affected by GALE depletion. By performing pathway enrichment and PPI clustering analysis on all differentially regulated N-linked glycoproteins following GALE depletion, we identified several recurrent clusters initially present in oncogenic RAS-transformed cells, including lysosome, glycoprotein metabolic process, extracellular matrix organization, cell adhesion, post-translational protein phosphorylation, wound healing, basigin interactions (from pathway analysis), as well as integrin signalling, Golgi associated vesicle biogenesis, and semaphorin-plexin signalling (from PPI analysis) (Fig. S4, K and L)^15^. We finally focused our analysis on N-linked glycoproteins that were hyposialylated following GALE depletion, common to both NIH 3T3 KRAS^G12V^ and PANC-1 cell models. This shared set represents a core group of GALE-dependent N-glycoproteins, each previously implicated in cancer progression, including BSG/CD147, CD276/B7-H3, CD44, ITGA6, ITGAV, ITGB1, MFGE8, MRC2, SLC3A2/CD98hc, and TFRC, for which we summarize the corresponding glycan-type and glycosite-specific changes (Fig. 4D). We further highlight one of these glycoproteins, namely CD44, a glycosylated cell surface receptor well-described to support the tumorigenic process, and summarize its putative glycan structures and glycosites in both GALE-replete and depleted contexts (Fig. 4E)^22^.

**Fig. 4.**
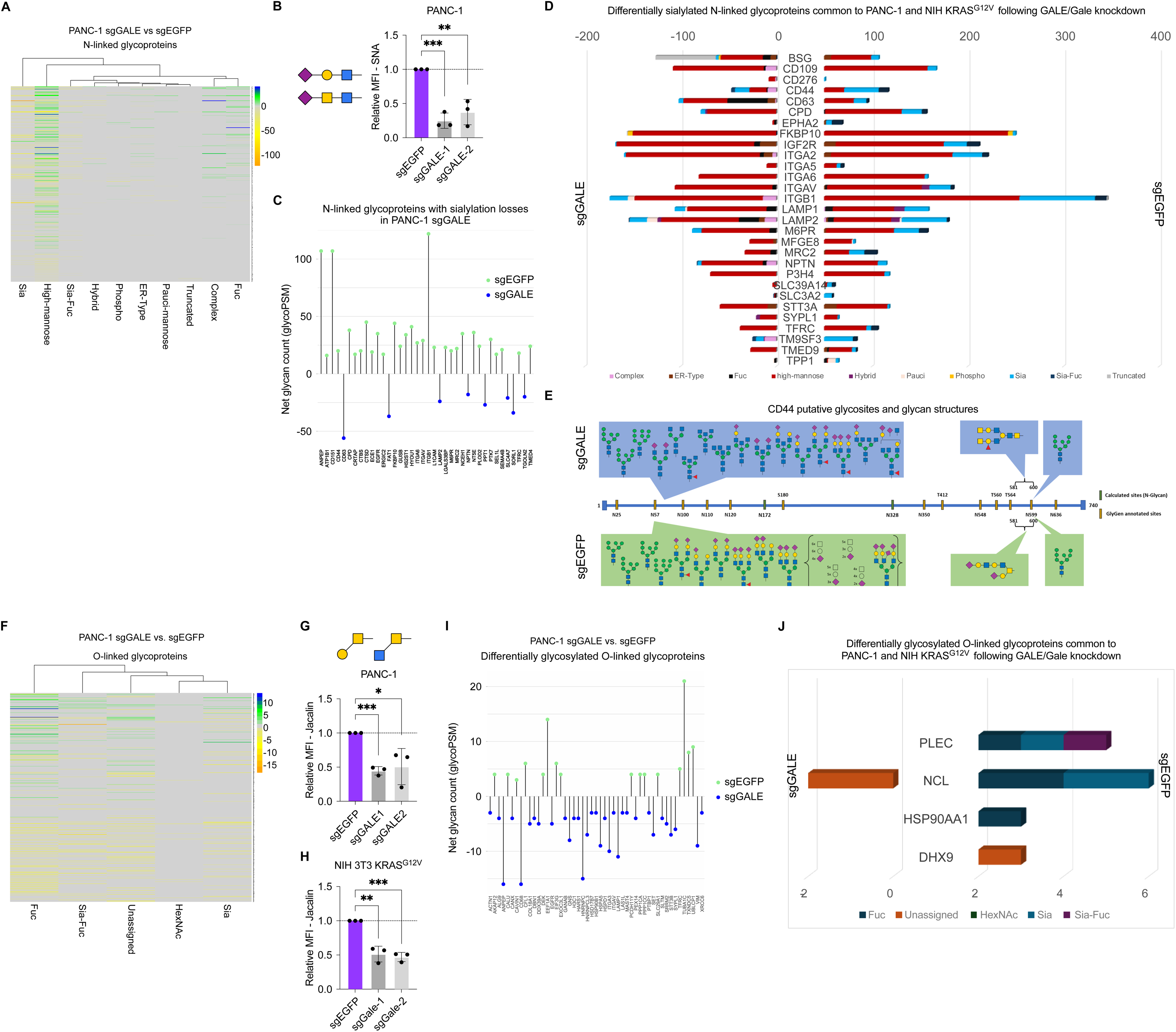
GALE mediates glycoproteome remodelling in oncogenic RAS-transformed cells. (A) Differential glycan analysis of N-linked glycoproteins comparing sgGALE versus sgEGFP PANC-1 cells, obtained via SAX-ERLIC enrichment method and tandem mass spectrometry. Counts represent glycopeptide-spectrum matches (glycoPSM). Sia represents sialylation; Sia-Fuc represents Sialyl-fucosylated. (B) SNA-fluorescein labelling of sgGALE-1, sgGALE-2, and sgEGFP PANC-1 cells. Relative MFI (mean fluorescence intensity) = [MFI_with lectin_-MFI_without lectin_]/MFI_without lectin_ normalized to the control from each biological replicate. (C) Net glycan changes (glycoPSM) across all glycan types for most differentially glycosylated N-linked glycoproteins (based on total absolute glycoPSM) in sgGALE PANC-1 relative to sgEGFP. (D) Differentially sialylated N-linked glycoproteins common to both PANC-1 and NIH 3T3 KRAS^G12V^ following GALE/Gale knockdown. The histogram shows the number of glycoPSM for each glycan type in sgGALE or sgEGFP PANC-1 cells. (E) Schematic showing the glycoprotein CD44 and theoretically mapped glycan structures and glycosites in sgGALE and sgEGFP PANC-1 cells. (F) Differential glycan analysis of O-linked glycoproteins comparing sgGALE versus sgEGFP PANC-1 cells, obtained as in (A). Fuc represents fucosylated; HexNAc represents single O-linked HexNAc; Unassigned represents general O-linked glycan. Jacalin-fluorescein labelling of (G) sgGALE-1, sgGALE-2, and sgEGFP PANC-1 cells, and (H) sgGale-1, sgGale-2, and sgEGFP NIH 3T3 KRAS^G12V^ cells. Relative MFI calculated as in (B). (I) Net glycan changes (glycoPSM) across all glycan types for most differentially glycosylated O-linked glycoproteins (based on total absolute glycoPSM) in sgGALE PANC-1 relative to sgEGFP. (J) Differentially glycosylated O-linked glycoproteins common to both PANC-1 and NIH 3T3 KRAS^G12V^ following *GALE/Gale* knockdown. The histogram shows the number of glycoPSM for each glycan type in sgGALE or sgEGFP PANC-1 cells. Unless otherwise indicated, data represent mean ± SD of at least three independent experiments. Unpaired, two-tailed *t test*; \**P* < 0.05, \*\**P* < 0.01, \*\*\**P* < 0.001, \*\*\*\**P* < 0.0001.

We further identified 494 O-linked glycoproteins with altered glycosylation following GALE depletion in PANC-1 cells (Fig. 4F, Table S5). These changes were characterized by a broad reduction in different types of O-linked glycans, consistent with decreased Jacalin and VVL binding in GALE-depleted cells relative to controls (Fig. 4G and H, Fig. S5A). These findings suggest reduced abundance of exposed terminal GalNAc residues and specific O-linked core GalNAc glycans, again in line with the enzymatic function of GALE in supplying the nucleotide-sugar precursors required for these glycoforms. Among the altered O-linked glycoproteins, pathways and PPI interaction analysis included recurrent enrichment of Rho GTPase signalling (TUBA1C) and vesicle-mediated transport (TFRC), alongside less anticipated clusters related to ribonucleotide metabolic process (EGFR) and RNA metabolism (PPP1CA, PPP1CC) (Fig. 4I, Fig. S5, B and C). We also focused our analysis on four altered O-linked glycoproteins—Plectin (PLEC), Nucleolin (NCL), Hsp90α (HSP90AA1) and DExH-Box Helicase 9 (DHX9)— that were common to both PANC-1 and NIH 3T3 KRAS^G12V^ cell models following GALE depletion and summarize the glycan type and glycosite changes that occur for each of these glycoproteins (Fig. 4J, Fig. S5, D and E). Overall, our data provide a comprehensive, site-resolved catalogue of candidate downstream targets of GALE-dependent N- and O-linked glycosylation, across two orthogonal *KRAS*-mutant cell lines, and providing attractive starting points for future interrogation.

### GALE-dependent glycosylation is a cell-intrinsic determinant of oncogenic RAS transformation and contributes to tumorigenicity in vivo

We next asked whether GALE function might contribute to oncogenic RAS transformation. To address this, we assessed the ability of *Gale*-knockdown and control NIH 3T3 KRAS^G12V^ cell lines to grow in soft-agar and observed that *Gale* knockdown markedly suppressed colony formation (Fig. 5A). In two other *KRAS*-mutant cell lines, *GALE* knockdown also led to a general suppression of soft-agar colony formation (Fig. 5B and Fig. S6A)^23^. Interestingly, *Gale* knockdown resulted in no observable effects on proliferation in 2-D monolayer cultures, suggesting that the negative effect of GALE depletion on soft-agar colony formation might be attributed to impaired anchorage-independent growth rather than reduced proliferative capacity (Fig. S6, B and C). Consistently, analysis of publicly available genome-wide CRISPR/Cas-9 perturbation screens indicates that GALE is not a determinant of cell growth in 2-D monolayer culture across human cancer cell lines (Fig. S6D).

**Fig. 5.**
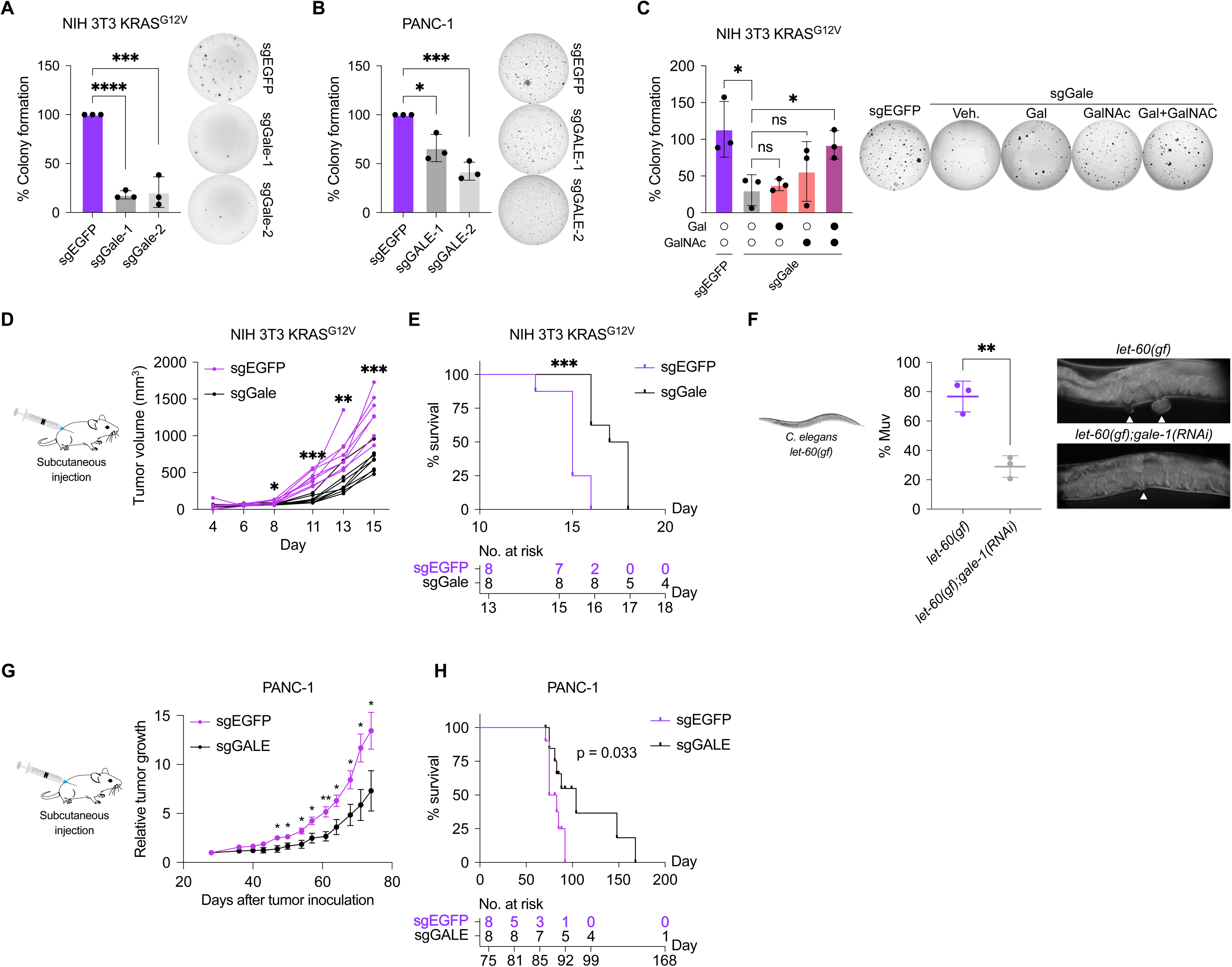
GALE contributes to oncogenic RAS transformation and tumorigenicity (A) Soft agar colony formation of CRISPR interference (CRISPRi)-mediated *Gale* knockdown (sgGale-1 and sgGale-2) and control (sgEGFP) NIH 3T3 cells expressing mutant KRAS^G12V^ (NIH 3T3 KRAS^G12V^. Colonies were imaged and quantified using ImageJ, and normalized to control. (B) Soft agar colony formation of CRISPR interference (CRISPRi)-mediated GALE knockdown (sgGALE-1 and sgGALE-2) and control (sgEGFP) PANC-1 cells. Colonies were imaged and quantified as in (A). (C) Soft agar colony formation of sgGale-1 and sgEGFP NIH 3T3 KRAS^G12V^ cells supplemented with 1 mM galactose (Gal) alone, 1 mM N-acetyl-galactosamine (GalNAc) alone or both in combination. Colonies were imaged and quantified as in (A). (D) sgGale-1 and sgEGFP NIH 3T3 KRAS^G12V^ cells were subcutaneously injected in the flank of NOD *SCID* gamma mice and tumor volumes were measured every 2-3 days. *n* = 8 for each cell line. (E) Survival of mice bearing tumors from (D) where the ethical endpoint of each mouse was considered an event. *n* = 8 for each cell line. (F) Percentage (%) Muv scored for *let-60(gf)* worms exposed to RNA interference (RNAi) targeting *gale-1* or controls. Each data point represents quantification from three independent experiments. (G) sgGALE (sgGALE-2) and sgEGFP PANC-1 cells were subcutaneously injected in the flank of NOD *SCID* gamma mice and tumor volumes were measured every 3-4 days. *n* = 8 for each cell line. (H) Survival of mice bearing tumors from (G) where the ethical endpoint of each mouse was considered an event. *n* = 8 for each cell line. Unless otherwise indicated, data represent mean ± SD of at least three independent experiments. Unpaired, two-tailed *t test*; \**P* < 0.05, \*\**P* < 0.01, \*\*\**P* < 0.001, \*\*\*\**P* < 0.0001. ns, not significant.

Intracellular UDP-galactose and UDP-GalNAc can be generated via GALE-dependent epimerization, but can also be supplied via alternative metabolic inputs, namely the Leloir pathway for galactose and salvage pathways for extracellular GalNAc. To ascertain whether the growth defects of *Gale*-knockdown cells under anchorage-independent conditions result from depletion of these nucleotide sugar pools, we supplemented culture media with galactose and/or GalNAc. Supplementation with either sugar alone showed no significant effects; however, combined supplementation markedly rescued colony formation in soft agar and increased tumor size in the chick chorio-allantoic membrane (CAM) assay (Fig. 5C and Fig. S6E). This functional rescue correlated with selective recovery of glycosylation: GalNAc supplementation restored VVL binding, whereas galactose supplementation restored SNA binding in GALE-deficient cells (Fig. S6F-H).

To investigate the contribution of GALE to oncogenic KRAS-mediated transformation in vivo, we performed subcutaneous injection of *Gale*-knockdown and control NIH 3T3 KRAS^G12V^ cells into immunodeficient mice. This revealed that GALE depletion substantially reduced tumor growth and prolonged host survival, as compared to controls (Fig. 5, D and E). We then asked whether this function of GALE is conserved in evolution. We previously employed the model organism *Caenorhabditis elegans* to provide a robust complementary system to study the in vivo functions of oncogenic RAS, whereby gain-of-function (*gf*) mutations in the *C. elegans Ras* ortholog *let-60* result in the development of multiple ectopic vulvae, termed the multivulva (Muv) phenotype^24^. Strikingly, RNA interference of *gale-1*, which encodes the *C. elegans* ortholog of mammalian *GALE*, suppressed the Muv phenotype of *let-60(gf)* worms, suggesting that the contribution of GALE function to oncogenic transformation is conserved across these biological contexts (Fig. 5F).

To determine whether the requirement for GALE is maintained in bona fide KRAS-mutant cancer cells, we evaluated GALE loss in PDAC tumorigenic models. In the CAM assay, *GALE* knockdown reduced tumor growth derived from PANC-1 and MIA PaCa-2 cells (Fig. S6, I and J). We then further validated these findings in a murine xenograft setting using PANC-1 cells, where GALE depletion led to a significant reduction in tumor progression and a concomitant survival benefit (Fig. 5, G and H). Together, these data demonstrate that GALE is a tumor cell-intrinsic determinant of oncogenic RAS-mediated transformation and tumor growth, extending beyond engineered RAS systems to established clinically relevant pancreatic cancer xenograft models.

### GALE is clinically relevant in pancreatic adenocarcinoma and other tumor entities

Next, we further investigated the potential clinical relevance of GALE in human PDAC. We found that *GALE* was overexpressed in tumor tissue from multiple orthogonal PDAC cohorts relative to non-tumor tissue, and that its expression was largely confined to malignant cells within the heterogenous tumor microenvironment (Fig. 6, A and B, Fig. S7A)^25–28^. Re-analysis of an institutional PDAC cohort from our previous study also showed that *GALE* was overexpressed in pancreatic intraepithelial neoplasia (PanIN) lesions, relative to normal tissue samples, suggesting *GALE* induction to be an early event in PDAC progression, consistent with our evidence that oncogenic RAS transformation leads to GALE upregulation (Fig. S7B)^29^. In concordance with transcript levels, GALE peptide abundance was similarly upregulated in PDAC relative to non-tumor tissue (Fig. 6C). To independently verify GALE expression in human PDAC, we analyzed a tissue microarray (TMA) comprising normal pancreas, primary tumors, and lymph node metastases using immunohistochemistry (IHC). GALE levels were significantly elevated in both primary and metastatic lesions compared to normal pancreatic tissue (Fig. 6D). Finally, higher expression of *GALE* correlated with shorter overall survival in patients with PDAC (Fig. 6E). These findings indicate a potential link between GALE levels and adverse clinical outcomes, and along with our earlier functional data, suggest that GALE might contribute to human PDAC development

**Fig. 6.**
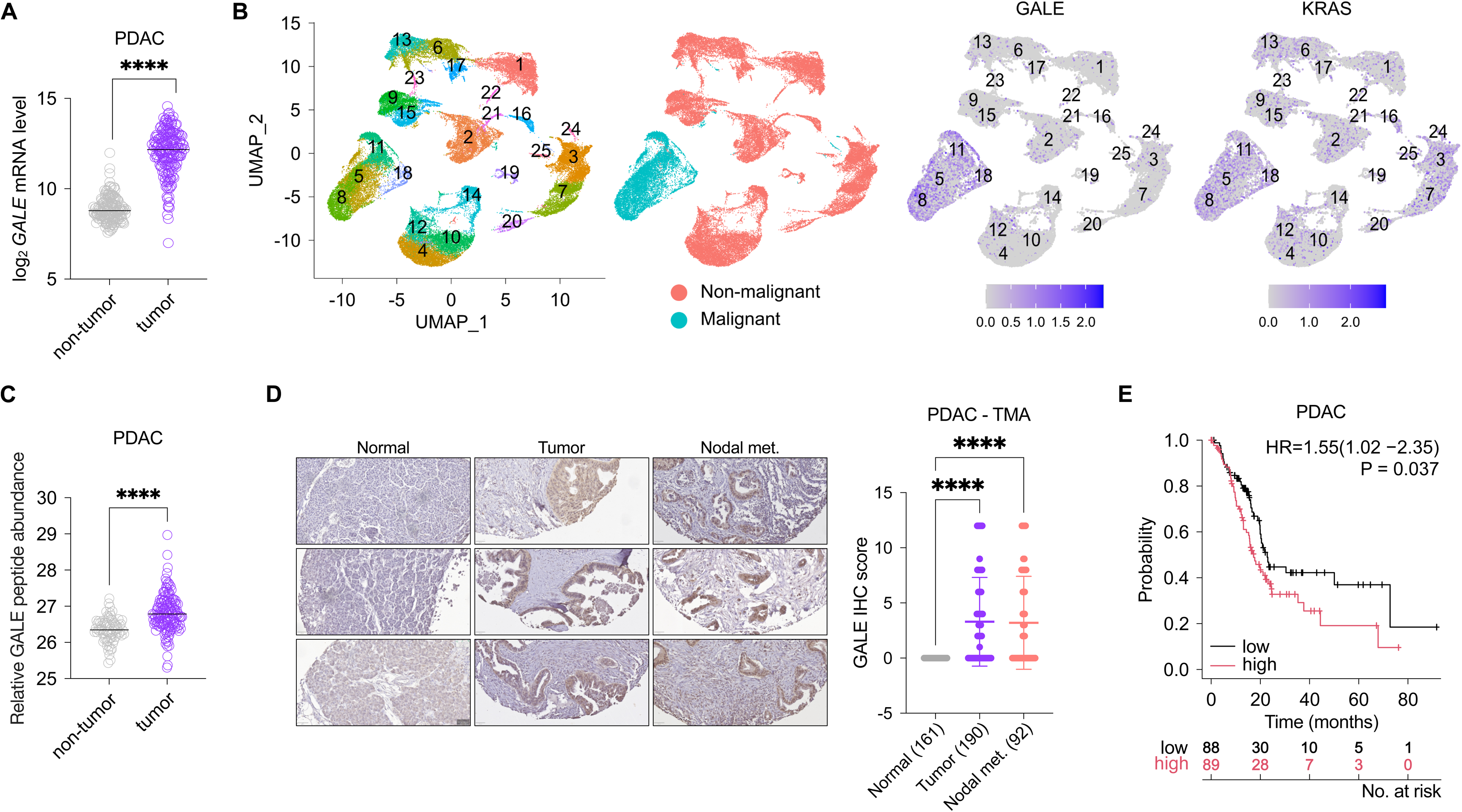
GALE expression is clinically relevant in PDAC. (A) *GALE* mRNA levels in PDAC samples from TCGA, in comparison to corresponding non-tumor tissue from TCGA and/or Genotype-Tissue Expression (GTEx) database. Data are analyzed using the UCSC Xena portal. (B) UMAP of single cell RNA sequencing data from 24 human PDAC samples showing malignant cell clusters, and expression of *GALE* and *KRAS*^25^. (C) GALE protein expression based on MS-proteomics in PDAC samples relative to non-tumor tissue from the Clinical Proteomic Tumor Analysis Consortium (CPTAC) study^53^. (D) Representative images of immunohistochemistry (IHC) of GALE in normal, tumor, and nodal metastasis samples from a human PDAC tissue microarray (TMA) and quantification using the Immunoreactive Scoring (IRS) system. Biologically independent samples for each group are indicated^29^. (E) Kaplan-Meier survival estimates of PDAC patients stratified by median *GALE* mRNA levels in the TCGA cohort. Data are analyzed using the KM-Plotter platform^54^. Unless otherwise indicated, data represent mean ± SD of at least three independent experiments. Unpaired, two-tailed *t test*; \**P* < 0.05, \*\**P* < 0.01, \*\*\**P* < 0.001, \*\*\*\**P* < 0.0001. ns, not significant.

We next asked whether the clinical relevance of *GALE* expression extends to other tumor entities. An analysis of tumor tissue data from The Cancer Genome Atlas (TCGA) in comparison to non-tumor tissue data from TCGA and/or The Genotype-Tissue Expression (GTEx) database revealed that *GALE* expression was significantly elevated across multiple tumor entities including non-small cell lung cancer, colon and rectal adenocarcinoma, skin cutaneous melanoma, breast invasive carcinoma, low-grade glioma, glioblastoma, neuroblastoma, thyroid cancer, diffuse large B-cell lymphoma, urothelial bladder carcinoma, cervical squamous cell carcinoma, ovarian serous cystadenocarcinoma, uterine corpus endometrial carcinoma, and uterine carcinosarcoma (Fig. 7A). Moreover, *GALE* expression was significantly associated with poorer outcome in colon and rectal adenocarcinoma, gastric adenocarcinoma, liver hepatocellular carcinoma, glioma, multiple myeloma, ovarian carcinoma, and sarcoma (Fig. 7B). Overall, these data support the potential clinical relevance of GALE not only for *KRAS*-mutant PDAC, but also for other human cancers.

**Fig. 7.**
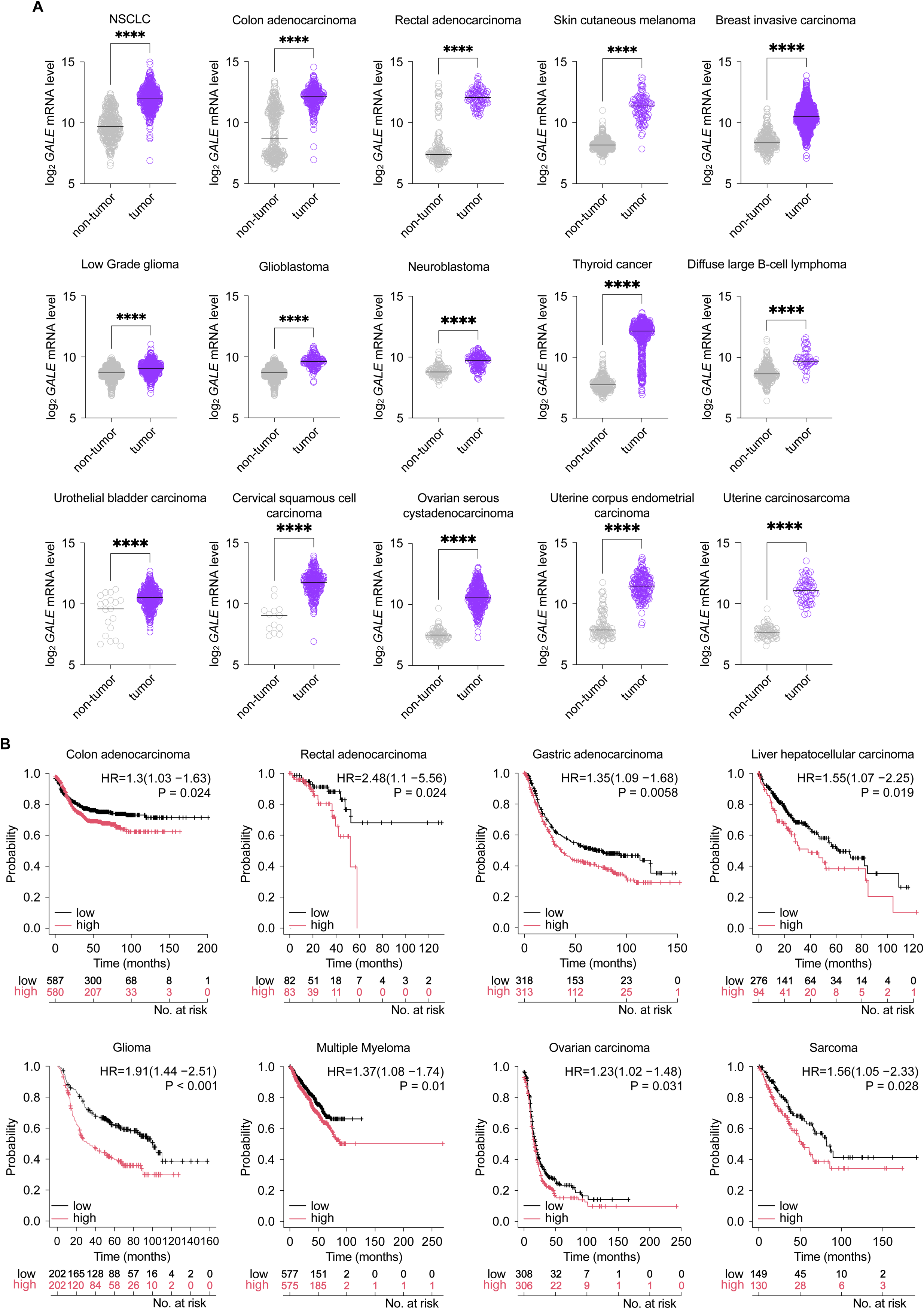
*GALE* expression is clinically relevant across multiple human cancers. (A) *GALE* mRNA levels in tumor samples from TCGA including non-small cell lung cancer, colon and rectal adenocarcinoma, skin cutaneous melanoma, breast invasive carcinoma, low-grade glioma, glioblastoma, neuroblastoma, thyroid cancer, diffuse large B-cell lymphoma, urothelial bladder carcinoma, cervical squamous cell carcinoma, ovarian serous cystadenocarcinoma, uterine corpus endometrial carcinoma, and uterine carcinosarcoma, in comparison to corresponding non-tumor tissue from TCGA and/or Genotype-Tissue Expression (GTEx) database. (B) Kaplan-Meier survival estimates of colon and rectal adenocarcinoma, gastric adenocarcinoma, liver hepatocellular carcinoma, glioma, multiple myeloma, ovarian carcinoma, and sarcoma patients stratified by median *GALE* mRNA levels (except for liver hepatocellular carcinoma, where cut-off was first-quartile) in the corresponding TCGA cohort. Data are analyzed using the KM-Plotter platform^54^. Unpaired, two-tailed *t test*; \**P* < 0.05, \*\**P* < 0.01, \*\*\**P* < 0.001, \*\*\*\**P* < 0.0001.

## Discussion

Aberrant glycosylation is a hallmark of cancer, yet how oncogenic RAS transformation reshapes the glycosylation landscape is incompletely understood. To address this, we employed a reductionist approach, namely the isogenic NIH 3T3 model with or without KRAS^G12V^ expression, which provides a robust model of transformation and tumorigenicity without the confounding effects of additional oncogenic lesions typically required to immortalize primary human cells, or tissue-specific lineage constraints. Using SAX-ERLIC enrichment followed by intact glycopeptide analysis, we show that oncogenic KRAS induces selective remodelling of the glycoproteome. The dominant changes to glycomic composition in KRAS-transformed cells were enrichment of sialylated and sialyl-fucosylated N-glycans, coupled with depletion of complex, hybrid, and high-mannose-glycans. These glycan alterations occurred on proteins with established roles in transformation and tumorigenicity, including mediators of cell-cell adhesion, extracellular matrix (ECM) remodelling, semaphorin-plexin signalling and integrin function, suggesting that glycosylation provides an additional regulatory layer within these pathways.

Earlier efforts to catalogue oncogenic RAS-regulated glycoproteins relied on hydrazide chemistry coupled to glycan release, generating protein-centric datasets biased towards N-linked glycoproteins and lacking site-specific glycopeptide information^30,31^. More recently, a comparative study using cell-surface capture combined with activated ion electron transfer dissociation (AI-ETD) resolved oncogene-specific glycoform patterns across multiple transformed cell lines. The study reported fewer high-mannose and sialylated N-linked glycopeptides, along with increased complex/hybrid-type glycopeptides in KRAS-mutant cells. Although we also observed an overall reduction in high-mannose glycans, these changes were bi-directional, with enrichment on some glycoproteins and depletion on others. Moreover, in contrast to that study, sialylation represented the predominantly enriched modification among N-linked glycoproteins after oncogenic KRAS transformation, whereas complex and hybrid glycans were significantly decreased. These differences may reflect methodological variation, including the use of ConA lectin enrichment and hydrophilic interaction liquid chromatography (HILIC), which preferentially recover oligomannose and hybrid glycans. Our findings align instead with other reports that mutant KRAS increases sialylation in murine tumor models, and with the broader literature identifying hypersialylation as a common feature of solid tumors, including PDAC^32–34^. In this regard, our data support the concept that oncogenic transformation contributes to the establishment of a tumor-associated “glycocode”. Whereas transcriptomic surveys and lectin-based profiling have been instrumental in defining cancer-associated glycosylation programmes, our study adds a site-resolved, intact glycoproteomic map of oncogenic KRAS-directed glycan remodelling^35^. Moreover, while most glycoproteomic studies in cancer have been optimized for investigating N-linked glycosylation, the O-linked glycoproteome remains comparatively underexplored (CPTAC study) owing to broader analytical constraints, including the absence of a consensus sequence for O-linked glycosite prediction, and a lack of enzymes that can release all types of O-linked glycans^36^. Yet, given that O-linked glycosylation is one of the most common and diverse types of glycosylation with essential roles in physiological and pathological processes, resolving these structures would undoubtedly lead to a more comprehensive understanding of their contribution to malignant phenotypes. Here, we found that O-linked glycoproteins differentially regulated by oncogenic KRAS more frequently exhibited enrichment of O-linked glycans across multiple glycan classes, and were associated with Rho GTPase signalling, actin cytoskeleton organization, collagen formation, and cell-cell communication, among other pathways and interaction networks. Interestingly, a subset of these O-linked glycoproteins are known nuclear proteins, including those involved in mRNA processing (HNRNPU, HNRNPA2B1, ILF3, HDGF; Fig. 1). While the role of intranuclear glycoproteins has been contentious, a recent report provided compelling evidence for a functional role of nuclear glycosylation, specifically showing that O-linked glycosylation of RNA-binding proteins was crucial for RNA processing^37^. Defining which of O-linked glycoproteins and O-glycosites functionally contribute to tumorigenicity, immune evasion, or therapeutic resistance will be an important area for future investigation. Thus, to our knowledge, this study provides the most comprehensive inventory of the oncogenic RAS-modulated glycoproteome to date, integrating quantitative glycoprotein abundance with site-resolved data on glycan composition and occupancy across both N-linked and O-linked glycoproteins.

Using a multiomics approach on our isogenic KRAS-transformed model, we discovered GALE as a transcriptional target of oncogenic RAS signalling. Across engineered and established KRAS-mutant cell models, we further demonstrate that GALE induction is mediated by MAPK signalling and the transcription factor MYC. Given that GALE converts glucose-derived nucleotide sugars into UDP-galactose and UDP-GalNAc, we hypothesized that GALE loss would preferentially affect galactose and GalNAc-containing glycans. In support of our hypothesis, Gale/GALE depletion in two distinct KRAS-mutant cell models caused a broad reduction in O-linked glycosylation, which is initiated by GalNAc, as well as decreased sialylation of N-linked glycans, in line with the fact that sialic acids are predominantly added to terminal galactose, GalNAc, or sialic acid residues, and echoing previous observations^38^. Nearly half of the N-linked glycoproteins modified during oncogenic KRAS transformation were also altered after GALE depletion, suggesting that GALE accounts for a substantial fraction of oncogenic KRAS-driven glycoproteome remodelling. By overlapping glycoproteomic data from both GALE-depleted KRAS-mutant models (NIH 3T3 KRAS^G12V^ and PANC-1), we define a site-resolved catalogue of N- and O-linked glycoproteins whose glycosylation status are uniformly GALE-dependent. Although hypersialylation of tumors has often been attributed to increased expression of sialyltransferases such as ST6GAL1 and ST3GAL1, evidence directly linking this process to oncogenic RAS signalling remains limited^32^. Our unbiased analyses instead identify GALE-dependent nucleotide-sugar precursor supply as a key mechanism by which oncogenic RAS reshapes the glycan landscape, including hypersialylation. These findings support a model in which RAS-MAPK-MYC signalling upregulates GALE, increasing UDP-galactose and UDP-GalNAc availability and thereby selectively privileging specific glycoforms over others. In this framework, altered glycosylation during oncogenic transformation emerges from the coordinated modulation of metabolic precursor supply, nucleotide sugar interconversion, and glycosyltransferase activity, extending previous work showing that mutant KRAS alters glycosylation through enhanced flux into the HBP. Further, the recurrent identification of specific cancer-associated glycoproteins, including CD44 and ITGB1, and associated networks, such as integrin signalling, cell-cell adhesion, and ECM organization, in our dataset and others suggests that oncogenic KRAS does not passively increase global glycosylation but instead establishes a prioritized glycoprotein programme that may support discrete tumorigenic functions^39,40^.

These findings may have important therapeutic implications, as GALE-dependent nucleotide-sugar interconversion may represent a mechanistic bottleneck in oncogene-driven glycoproteome remodelling. In support of this idea, GALE knockdown impaired anchorage⍰independent growth of oncogenic KRAS⍰transformed cells and suppressed tumor growth in murine xenograft models, positioning GALE as a potential therapeutic vulnerability in RAS⍰mutant cancers. Notably, GALE depletion had no observable effects on proliferation under 2⍰D culture conditions, suggesting that GALE inhibition may preferentially target malignant growth states while sparing non-tumor tissues. This possibility is further supported by genetic evidence indicating that GALE is essential during *Drosophila* embryonic development but has less clearly defined roles in adult tissues, and by the observation that inherited GALE mutations in humans cause rare metabolic disorders that typically present in early childhood and often attenuate during adolescence and adulthood^41,42^. Additionally, GALE is overexpressed not only in *KRAS*⍰mutant cancers including PDAC, but also across tumor types in which *KRAS* mutations are less prevalent, suggesting broader clinical relevance. Our anchorage-independence and xenograft studies support a tumor cell-intrinsic role for GALE, as these models are devoid of an intact immune compartment. Along with our glycoproteomic data, these findings substantiate a role of GALE in directly sustaining a tumor-supporting glycoproteome independently of immune interactions. However, this also represents an important limitation that needs to be addressed in the future, as the contribution of GALE-dependent glycosylation to anti-tumor immunity remains unresolved. This question is particularly relevant because tumors frequently exhibit hypersialylation, which can mask tumor antigens and engage inhibitory Siglec receptors on immune cells. Indeed, the sialic acid–Siglec axis functions as a glyco-immune checkpoint that can suppress natural killer (NK) cell cytotoxicity, macrophage phagocytosis and T⍰cell activation^33,43,44^. Our observation that GALE knockdown reduces sialylation raises the possibility that GALE inhibition could decrease sialylated TACAs, including sialyl⍰Tn and sialyl⍰Lewis structures, as well as broader classes of tumour-associated Siglec ligands, thereby limiting Siglec engagement and enhancing immune recognition. In this context, GALE inhibition may complement emerging strategies targeting glyco-immune checkpoints, including Siglec blockade, desialylating biologics such as antibody–sialidase conjugates, and immune checkpoint inhibitors^45–47^. These possibilities remain speculative and would need to be evaluated in immunocompetent syngeneic models.

Whether GALE also contributes to metastasis, invasion, or therapy resistance remains unknown. Moreover, although our study used genetic depletion, pharmacological GALE inhibition has not yet been evaluated in this setting. Recent identification of a ligandable pocket near the human GALE active site, coupled with the development of covalent and non-covalent inhibitors, supports the feasibility of directly targeting this enzyme^48^. Similarly, 4-fluorosamine was shown to inhibit GALE activity, deplete UDP-GalNAc pools and reduce chondroitin sulfate levels^49^. Together, these studies support GALE as a chemically tractable enzyme and provide a foundation for future inhibitor development.

Collectively, our work provides a site-resolved framework for understanding how oncogenic RAS reshapes the glycoproteome to support malignant transformation and identifies GALE-dependent nucleotide-sugar interconversion as a central mechanism through which this occurs. More broadly, we establish sugar-precursor metabolism as a regulatory layer of oncogenic signalling and nominate GALE as a tractable vulnerability in RAS-driven and glycosylation-dependent cancers.

## Materials and Methods

### Cell culture

NIH 3T3 cells expressing mutant KRAS^G12V^ and control murine stem cell virus (MSCV) vectors were generated and cultured as previously described^50^. BJ cells expressing human telomerase (hTERT), SV40 large T (LT) and mutant *HRAS^G12V^* (hTERT/LT/HRAS^G12V^) and control hTERT/LT were a kind gift from William C. Hahn (Dana-Farber Cancer Institute, Boston, U. S. A). H460 cells were a kind gift from Julian Downward (Francis Crick Institute, London, UK). PANC-1, MIA PaCa-2, CFPAC1 and KPC cells were a kind gift from Barbara Grüner (German Cancer Consortium DKTK, Essen, Germany). KP cells were a kind gift from Paolo Ceppi (University of Southern Denmark, Odense, Denmark). PaTu-8988T and HEK293T cells were purchased from American Type Culture Collection (Rockville, MD). Unless otherwise specified, all cell lines were maintained in DMEM (Gibco) supplemented with 10% fetal calf serum (FCS; Sigma Aldrich). All cells were kept at 37 °C with 5% CO2. All cell lines were routinely confirmed to be mycoplasma-free using Venor®GeM Classic kit (Minerva Biolabs, Berlin, Germany). All human cell lines were authenticated by STR-profiling (Genomics and Transcriptomics Laboratory, Heinrich-Heine University, Germany).

### Lectin assay

Lectin assays were performed using fluorescein-conjugated lectins (SNA, Jacalin, WGA, VVL; Vector Laboratories) or biotin-conjugated lectins (Lectin kits I-III; Vector Laboratories). Concentrations used were according to manufacturer’s guidelines or determined by using different concentrations and establishing the right binding conditions. Cells were trypsinized, centrifuged, washed with PBS and 1*10^6 cells were resuspended in 1% BSA in PBS with appropriate amounts of lectin and incubated for 30-60 minutes on a rotator at 4°C in the dark. After washing two times with 1% BSA in PBS, the cells were either resuspended in 1 mL 1% BSA in PBS or, when previously incubated with biotin-conjugated lectins, incubated in a 1:10,000 dilution of a secondary streptavidin-FITC (Invitrogen) antibody. Afterwards, the cells were also washed two times with 1% BSA in PBS and resuspended again to form a single cell suspension. Analysis was done by flow cytometry using a BD FACSDiva. At least 10,000 events were recorded for each replicate and analyzed using FlowJo software.

Full Materials and Methods available in Supplemental information

## Acknowledgements

The authors gratefully acknowledge Stefanie Lichtenberg and Katharina Raba from the Core Facility Flow Cytometry of the Medical Faculty, Heinrich Heine University Düsseldorf, Germany for their support and assistance in this work. The authors also gratefully acknowledge Sandra Biskup and Olga Felda from the Institute of Pathology, University Hospital Düsseldorf, for their support and technical assistance. Research in the lab of J.K.M.L was supported by the Deutsche Forschungsgemeinschaft (Walter Benjamin Fellowship no. LI3844/1-1), Dr. Rolf M. Schwiete Foundation (grant no. 2023-029), Brigitte and Dr. Konstanze Wegener Foundation (grant no. 138: A2024 II # 9), the Research Commission of the Medical Faculty, Heinrich Heine University Düsseldorf (grant no. 2022-12), and the Volkswagen Foundation (grant no. Az. 9D177). N.S.C was supported by Aarhus Universitets Forskningsfond (Recruiting Grant AUFF-E-2022-7-10). Research in the lab of G.F. was supported by the Deutsche Forschungsgemeinschaft (FL865/2-1) and the Research Commission of the Medical Faculty, Heinrich Heine University Düsseldorf (grant no. 2024-60). W.B.D. is the Canada Research Chair in Animal Models of Human Disease, and supported by a Project Grant from the Canadian Institutes of Health Research (PJT 165837). The results presented here are in part based upon data generated by the TCGA Research Network: https://www.cancer.gov/tcga.

## Author contributions

Conceptualization: J.K.M.L. and U.G.D. Methodology: J.K.M.L., U.G.D., M.D.D., N.S.C., I.E., G.F., P.S., G.R., G.L., W.B.D., D.K., M.R. Investigation: J.K.M.L., F.S., R.V., C.R.S., M.C., A.D., Q.L., C.B., S.C., R.L., Ru. L., B.Y., D.P., V.S., F.C., A.Y., S.A.S., W.T.K., L.H. Writing – Original Draft, J.K.M.L; Writing – Review & Editing: F.S., C.R.S., M.C., A.D., G.F., G.L., G.R., M.D.D., U.G.D, J.K.M.L. Funding Acquisition: J.K.M.L. and U.G.D.

---

**Fig. S1.**
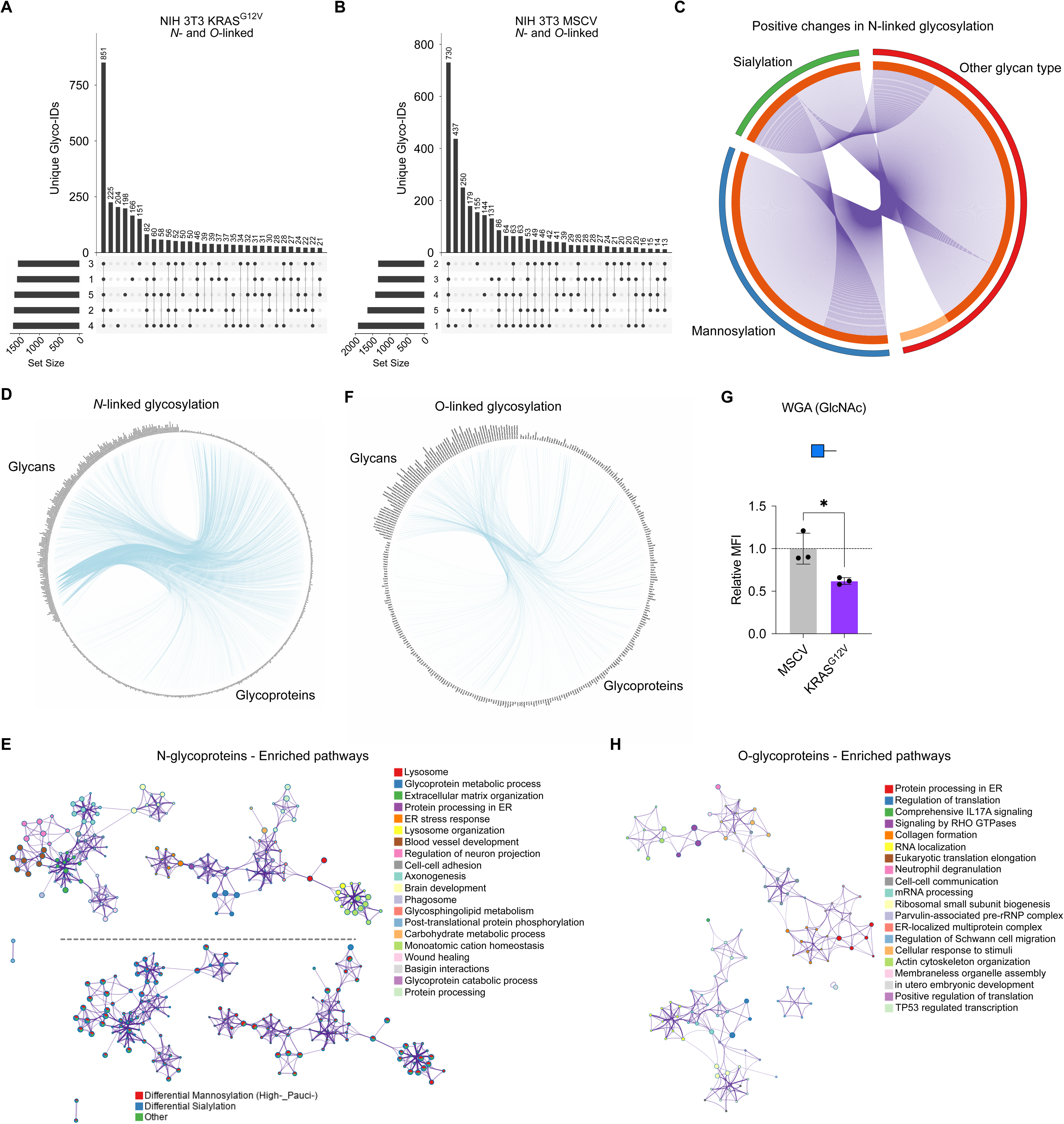
UpSet plots displaying N- and O-linked glycopeptide identification (Unique Glyco-IDs) in (A) NIH 3T3 KRAS^G12V^ and (B) NIH 3T3 MSCV cells that are unique to or shared across five independent replicates, including set sizes of each replicate. (C) Circos plot showing all N-linked glycoproteins with positive changes to any glycan type, and overlap between dominant glycan types (sialylation, high-/pauci-mannose) or other glycan type. Outside arc color identifies each glycan type by color. Inside arc represents glycoproteins containing associated glycan type; dark orange bars represent glycoproteins shared between datasets linked with purple lines, and light orange bars represent non-overlapping glycoproteins. (D) Chord diagram representing qualitative glycan-protein relationships of differentially altered N-linked glycoproteins comparing NIH 3T3 KRAS^G12V^ to NIH 3T3 MSCV cells. Glycan moieties are indexed to the top-left of the diagram and connected via chords to respective identified glycoproteins at the bottom-right. (E) Network shows pathway enrichment of all N-linked glycoproteins with positive changes to any glycan type. *top*, Each node is identified by color associated with pathway. Size of clusters is proportional to the number of input proteins included in the term. *bottom*, Each node displays the contribution of glycan type, including mannosylation or sialylation or other. (F) Chord diagram representing qualitative glycan-protein relationships of differentially altered O-linked glycoproteins comparing NIH 3T3 KRAS^G12V^ to NIH 3T3 MSCV cells, organized as in (D). (G) WGA-fluorescein labelling of NIH 3T3 KRAS^G12V^ and MSCV cells. Relative MFI (mean fluorescence intensity) = [MFI_with lectin_-MFI_without lectin_]/MFI_without lectin_ normalized to the control from each biological replicate. (H) Network shows pathway enrichment of all O-linked glycoproteins with positive changes to any glycan type. Each node is identified by color associated with pathway. Size of clusters is proportional to the number of input proteins included in the term. Unless otherwise indicated, data represent mean ± SD of at least three independent experiments. Unpaired, two-tailed *t test*; \**P* < 0.05, \*\**P* < 0.01, \*\*\**P* < 0.001, \*\*\*\**P* < 0.0001. ns, not significant.

**Fig. S2.**
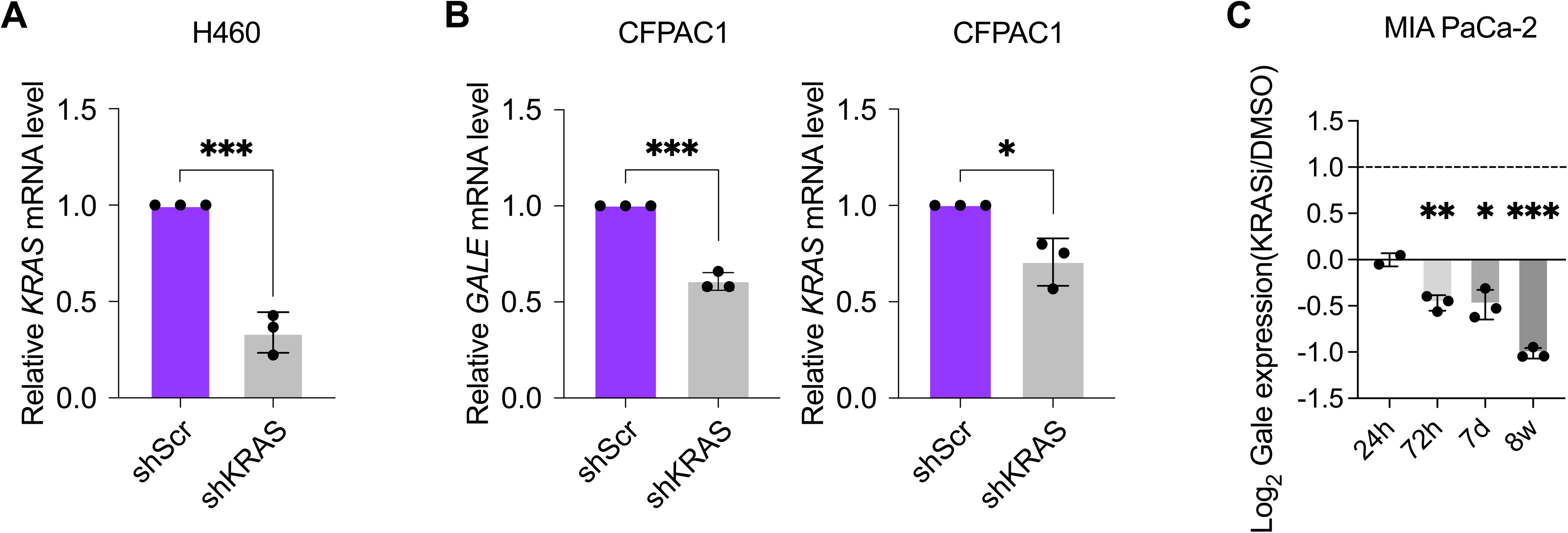
(A) Relative *KRAS* mRNA expression in shKRAS knockdown and scrambled (scr) control H460 cells, analyzed by qRT-PCR. (B) Relative *GALE* and *KRAS* mRNA expression in shKRAS knockdown and scrambled (scr) control CFPAC1 cells, analyzed by qRT-PCR. (C) Log_2_ *GALE* mRNA expression in MIA PaCa-2 cells upon treatment with 2 µM ARS-1620 (KRASi) for the indicated timepoints. Data were analyzed from https://manciaslab.shinyapps.io/KRASi/^55^. Unless otherwise indicated, data represent mean ± SD of at least three independent experiments. Unpaired, two-tailed *t test*; \**P* < 0.05, \*\**P* < 0.01, \*\*\**P* < 0.001, \*\*\*\**P* < 0.0001. ns, not significant.

**Fig. S3.**
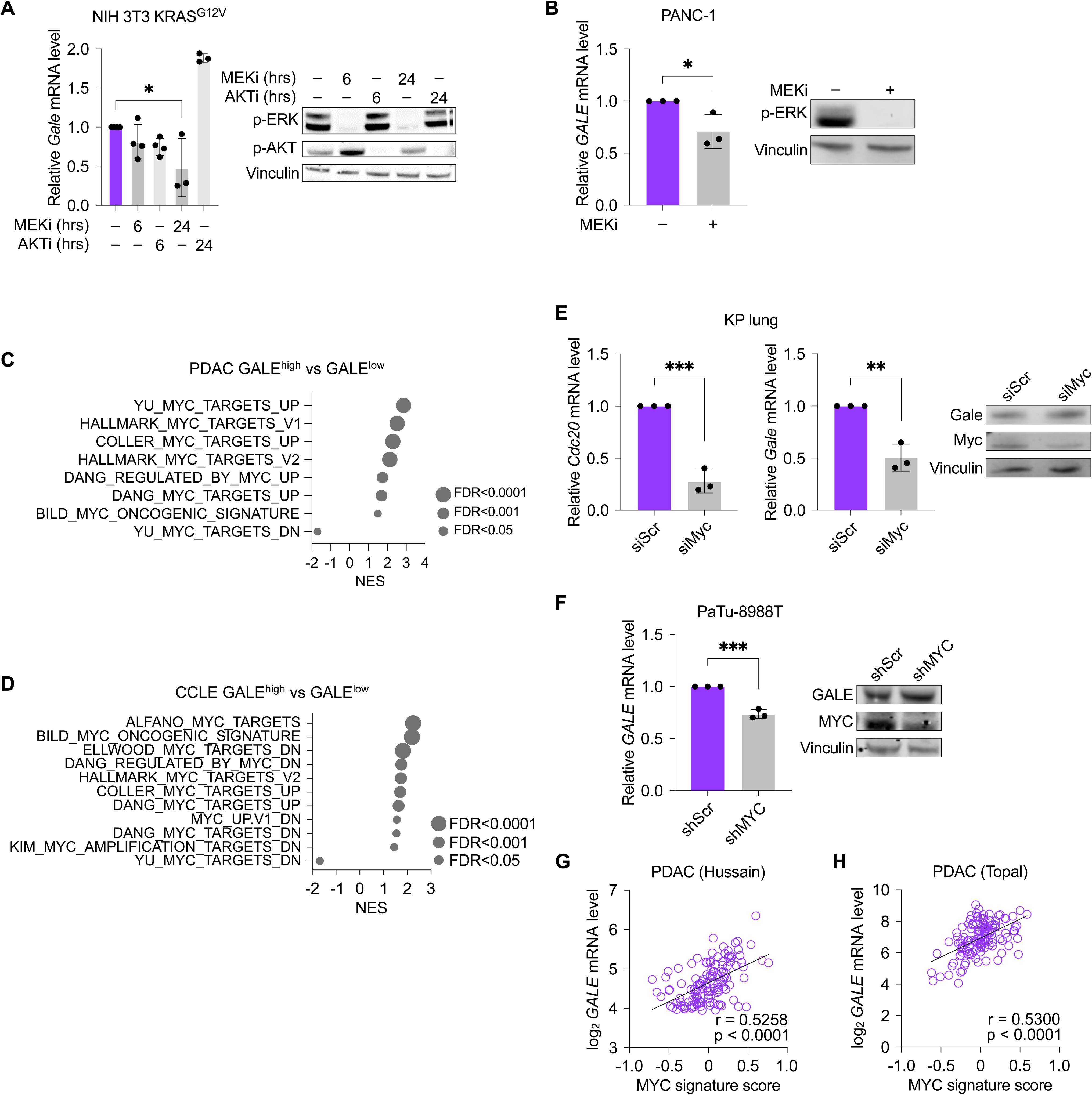
(A) Relative *Gale* mRNA expression analyzed by qRT-PCR, and expression of indicated proteins analyzed by Western Blot in NIH 3T3 cells expressing mutant KRAS^G12V^ (NIH 3T3 KRAS^G12V^) upon treatment with MEK inhibitor PD184352 (MEKi; 10 µM), AKT inhibitor MK2206 (AKTi ; 10 µM) or vehicle for the indicated times. (B) Relative *GALE* mRNA expression analyzed by qRT-PCR, and expression of p-ERK and Vinculin, analyzed by Western Blot in PANC-1 cells upon treatment with MEK inhibitor PD184352 (MEKi; 20 µM) or vehicle for 72 hours. Supervised gene set enrichment analysis (GSEA) using Human Molecular Signatures Database (MSigDB) MYC-related gene sets within “transcription factor targets (TFT)” performed on *GALE*-high versus GALE-low samples from (C) pancreatic ductal adenocarcinoma (PDAC) in The Cancer Genome Atlas (TCGA), and (D) cancer cell lines in the Cancer Cell Line Encyclopedia (CCLE) database. (E) Relative *Gale* and *Cdc20* mRNA expression analyzed by qRT-PCR, and protein expression of GALE, MYC, and Vinculin analyzed by Western Blot in KP cells transfected with siRNA against *Myc* (siMyc) or scrambled (siScr) control. (F) Relative *GALE* mRNA expression analyzed by qRT-PCR, and protein expression of GALE, MYC, and Vinculin analyzed by Western Blot in PaTu-8988T cells transduced with shRNA against *MYC* (shMYC) or scrambled (shScr) control. Correlation between MYC signature score and *GALE* expression in PDAC samples from (G) GSE62452 (Hussain cohort) and (H) GSE62165 (Topal cohort)^26,27^. Correlation was quantified using Spearman’s rank correlation coefficient. Data were analyzed via the R2 platform (http://r2.amc.nl)^21^. Unless otherwise indicated, data represent mean ± SD of at least three independent experiments. Unpaired, two-tailed *t test*; \**P* < 0.05, \*\**P* < 0.01, \*\*\**P* < 0.001, \*\*\*\**P* < 0.0001. ns, not significant.

**Fig. S4.**
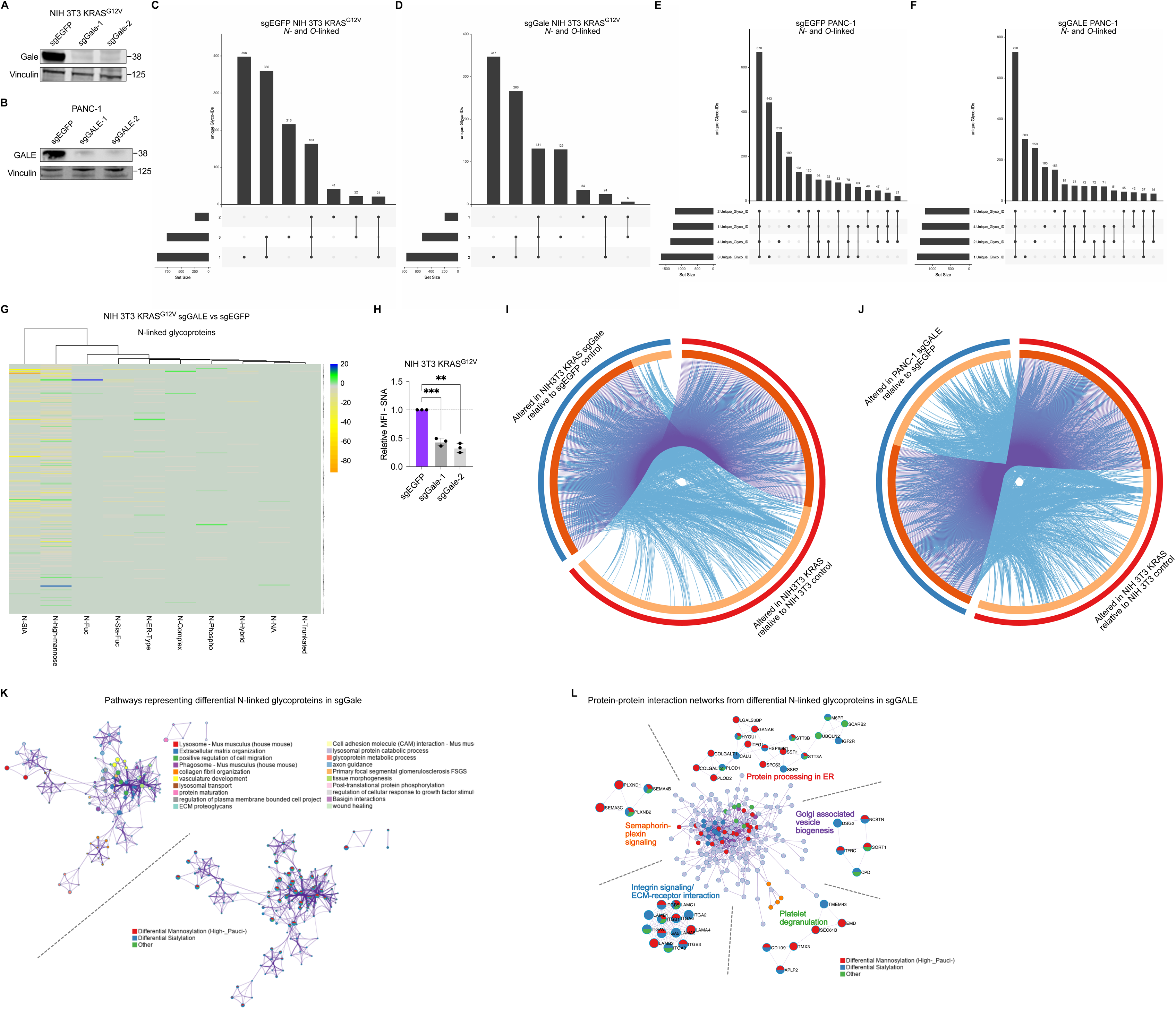
GALE and Vinculin expression in (A) sgGale-1, sgGale-2, and sgEGFP control NIH 3T3 KRAS^G12V^ cells and (B) sgGALE-1, sgGALE-2, and sgEGFP control PANC-1 cells. UpSet plots displaying N- and O-linked glycopeptide identification (Unique Glyco-IDs) in (C) sgEGFP control NIH 3T3 KRAS^G12V^ and (D) sgGale NIH 3T3 KRAS^G12V^ cells that are unique to or shared across three independent replicates, including set sizes of each replicate. UpSet plots displaying N- and O-linked glycopeptide identification (Unique Glyco-IDs) in (E) sgEGFP control PANC-1 and (F) sgGALE PANC-1 cells that are unique to or shared across four independent replicates, including set sizes of each replicate. (G) Circos plot showing overlap between N-linked glycoproteins differentially glycosylated in sgGale vs sgEGFP NIH 3T3 KRAS^G12V^, and those differentially glycosylated in NIH 3T3 KRAS^G12V^ vs MSCV cells. Outside arc color identifies glycoprotein dataset. Dark orange inside arc represents glycoproteins shared between datasets, linked by purple lines, and light orange arc represents non-overlapping glycoproteins. Blue lines connect glycoproteins with the same GO term. (H) SNA-fluorescein labelling of sgGale-1, sgGale-2, and sgEGFP NIH 3T3 KRAS^G12V^ cells. Relative MFI (mean fluorescence intensity) = [MFI_with lectin_-MFI_without lectin_]/MFI_without lectin_ normalized to the control from each biological replicate. (I) Circos plot showing overlap between N-linked glycoproteins differentially glycosylated in sgGale vs sgEGFP NIH 3T3 KRAS^G12V^ cells, and in NIH 3T3 KRAS^G12V^ vs MSCV cells. Outside arc color identifies glycoprotein dataset. Dark orange inside arc represents glycoproteins shared between datasets, linked by purple lines, and light orange arc represents non-overlapping glycoproteins. Blue lines connect glycoproteins with the same GO term. (J) Circos plot showing overlap between N-linked glycoproteins differentially glycosylated in sgGALE vs sgEGFP PANC-1 cells, and in NIH 3T3 KRAS^G12V^ vs MSCV cells. Outside arc color identifies glycoprotein dataset. Dark orange inside arc represents glycoproteins shared between datasets, linked by purple lines, and light orange arc represents non-overlapping glycoproteins. Blue lines connect glycoproteins with the same GO term. (K) Network shows pathway enrichment of differentially glycosylated N-linked glycoproteins in sgGale NIH 3T3 KRAS^G12V^ cells. *top*, Each node is identified by color associated with pathway. Size of clusters is proportional to the number of input proteins included in the term. *bottom*, Each node displays the contribution of glycan type, including mannosylation or sialylation or other. (L) Protein-protein interaction (PPI) analysis of differentially glycosylated N-linked glycoproteins in sgGALE PANC-1 cells, with the contribution of high- and pauci-mannosylated N-glycoproteins, sialylated (including Sia-Fuc), and other glycan type indicated. The center network represents the full interactome, and the surrounding networks indicated representative functions. Unless otherwise indicated, data represent mean ± SD of at least three independent experiments. Unpaired, two-tailed *t test*; \**P* < 0.05, \*\**P* < 0.01, \*\*\**P* < 0.001, \*\*\*\**P* < 0.0001. ns, not significant.

**Fig. S5.**
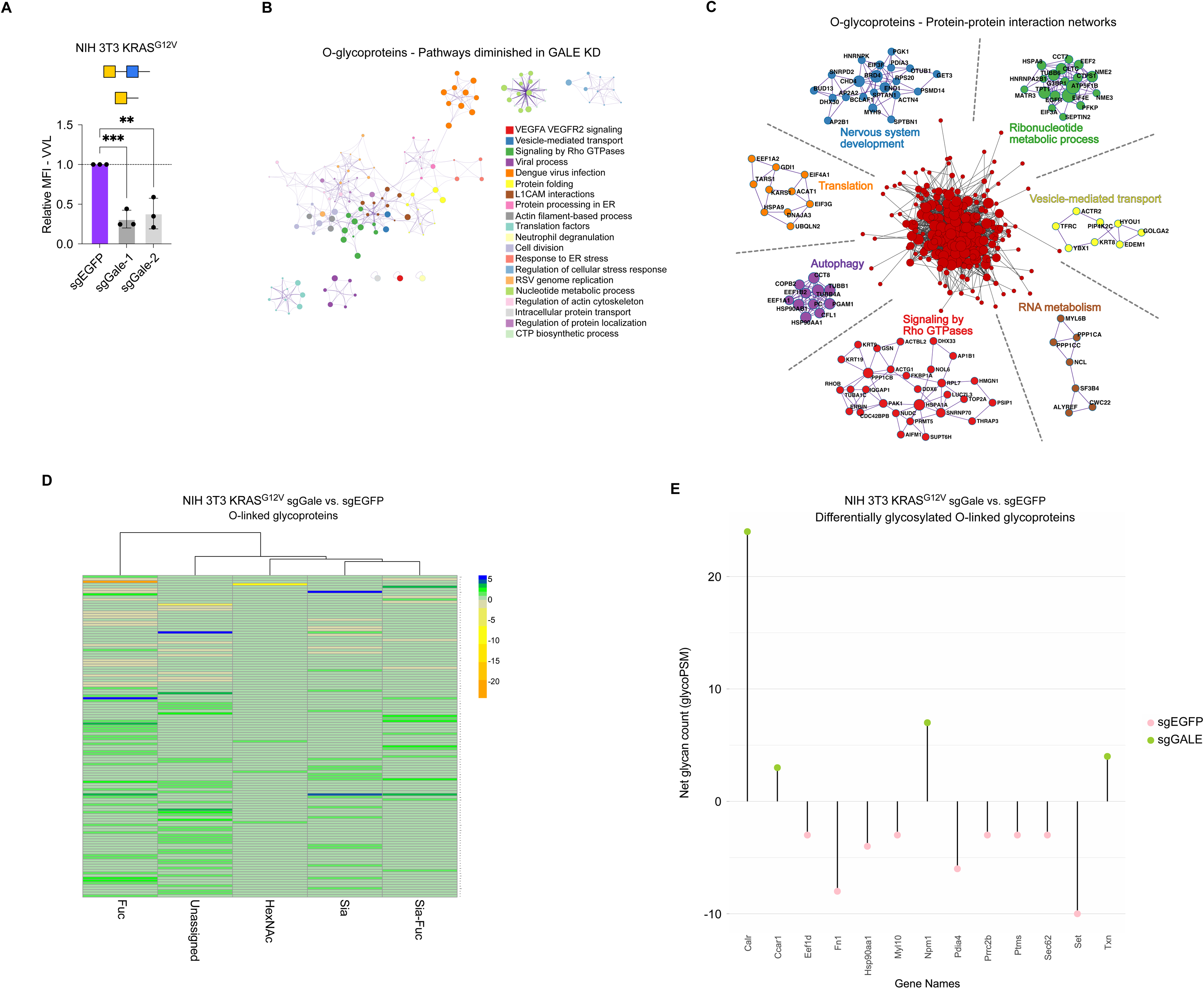
(A) VVL-fluorescein labelling of sgGale-1, sgGale-2, and sgEGFP NIH 3T3 KRAS^G12V^ cells. Relative MFI (mean fluorescence intensity) = [MFI_with lectin_-MFI_without lectin_]/MFI_without lectin_ normalized to the control from each biological replicate. (B) Network shows pathway enrichment of differentially glycosylated O-linked glycoproteins in sgGALE PANC-1 cells. Each node is identified by color associated with pathway. Size of clusters is proportional to the number of input proteins included in the term. (C) Protein-protein interaction (PPI) analysis of differentially glycosylated O-linked glycoproteins in sgGALE PANC-1 cells. The center network represents the full interactome, and the surrounding networks indicated representative functions. (D) Differential glycan analysis of O-linked glycoproteins comparing sgGale versus sgEGFP NIH 3T3 KRAS^G12V^ cells. Fuc represents fucosylated; HexNAc represents single O-linked HexNAc; Unassigned represents general O-linked glycan. (E) Net glycan changes (glycoPSM) across all glycan types for O-linked glycoproteins showing the most significant glycan losses (based on total absolute glycoPSM) in sgGale NIH 3T3 KRAS^G12V^ relative to sgEGFP. Unless otherwise indicated, data represent mean ± SD of at least three independent experiments. Unpaired, two-tailed *t test*; \**P* < 0.05, \*\**P* < 0.01, \*\*\**P* < 0.001, \*\*\*\**P* < 0.0001. ns, not significant.

**Fig. S6.**
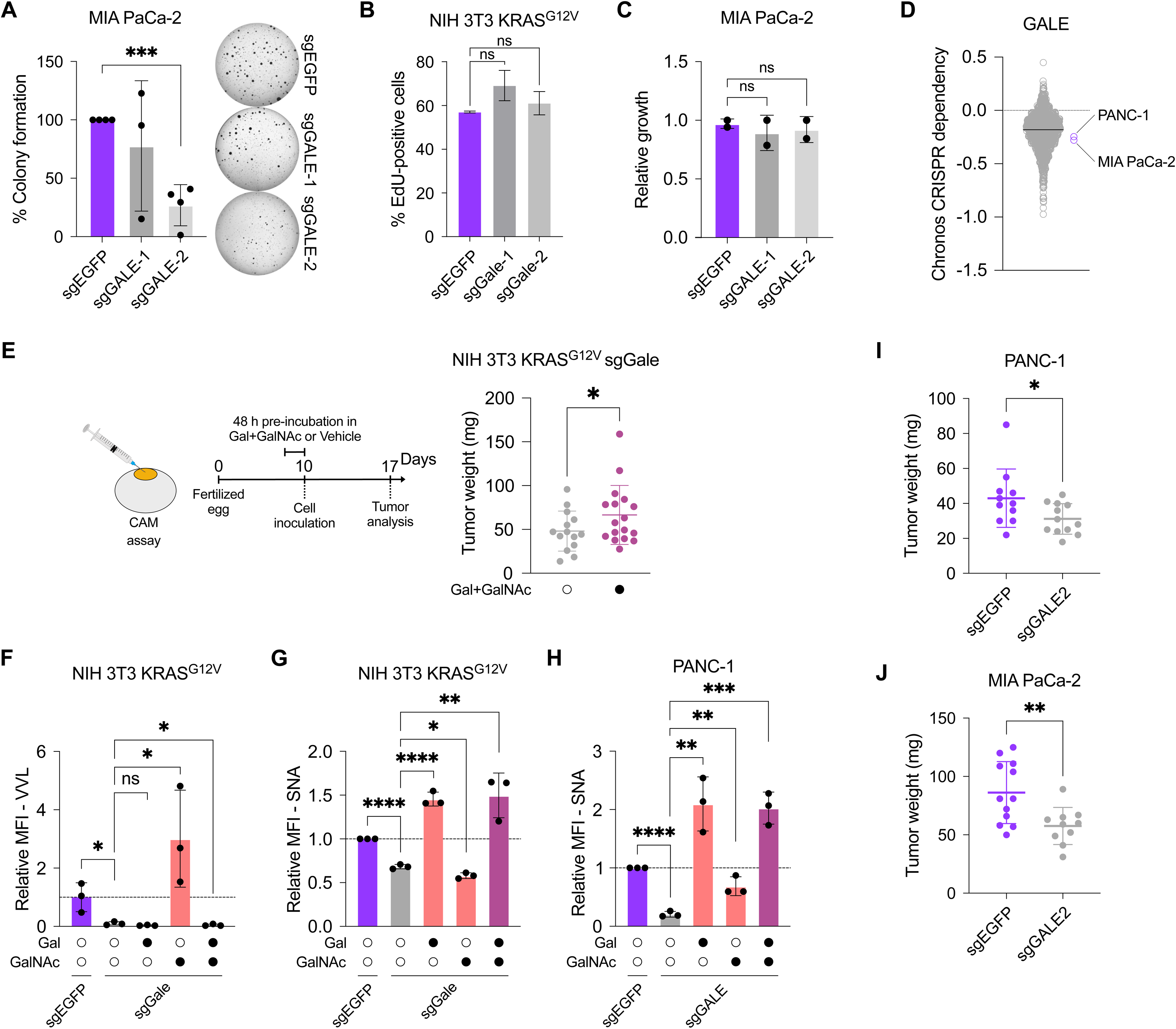
(A) Soft agar colony formation of CRISPR interference (CRISPRi)-mediated *GALE* knockdown (sgGALE-1 and sgGALE-2) and control (sgEGFP) MIA PaCa-2 cells. Colonies were imaged and quantified using ImageJ, and normalized to control. (B) Proliferation of CRISPR interference (CRISPRi)-mediated *Gale* knockdown (sgGale-1 and sgGale-2) and control (sgEGFP) NIH 3T3 KRAS^G12V^ cells, measured using EdU assay (*n=2*). (C) Relative growth of sgGALE-1 and sgGALE-2 and control sgEGFP MIA PaCa-2 cells, measured using Incucyte analysis (*n=2*). (D) Chronos CRISPR dependency scores for GALE in cancer cell lines from the DepMap consortium, with PANC-1 and MIA PaCa-2 highlighted. (E) sgGale (sgGale-1) NIH 3T3 KRAS^G12V^ cells with or without pre-incubation for 48 hours in standard culture media containing galactose and N-acetylgalactosamine (Gal+GalNAc) were inoculated onto the upper chorioallantoic membrane (CAM) of post-fertilized (d7) specific-pathogen-free chicken (SPF) eggs according to the indicated scheme, following which tumors were harvested and measured 7 days following inoculation. (F) VVL-fluorescein and (G) SNA-fluorescein labelling of sgEGFP and sgGale-1 (sgGale) NIH 3T3 KRAS^G12V^ cells with or without 48-hour supplementation with 1 mM Gal, 1 mM GalNAc, or both. (H) SNA-fluorescein labelling of sgEGFP and sgGALE-2 (sgGALE) PANC-1 cells with or without 48-hour supplementation with 1 mM Gal, 1 mM GalNAc, or both. The relative mean fluorescence intensity (rMFI= [MFI_with lectin_-MFI_without lectin_]/MFI_without lectin_) normalized to the corresponding control (sgEGFP) of each biological replicate. (I) sgEGFP (*n=11*) and sgGALE-2 (sgGALE; *n=12*) PANC-1 cells and (J) sgEGFP (*n=12*) and sgGALE-2 (sgGALE; *n=10*) MIA PaCa-2 cells were inoculated onto the CAM of post-fertilized (d7) SPF eggs, following which tumors were harvested and measured 7 days following inoculation. Unless otherwise indicated, data represent mean ± SD of at least three independent experiments. Unpaired, two-tailed *t test*; \**P* < 0.05, \*\**P* < 0.01, \*\*\**P* < 0.001, \*\*\*\**P* < 0.0001.

**Fig. S7.**
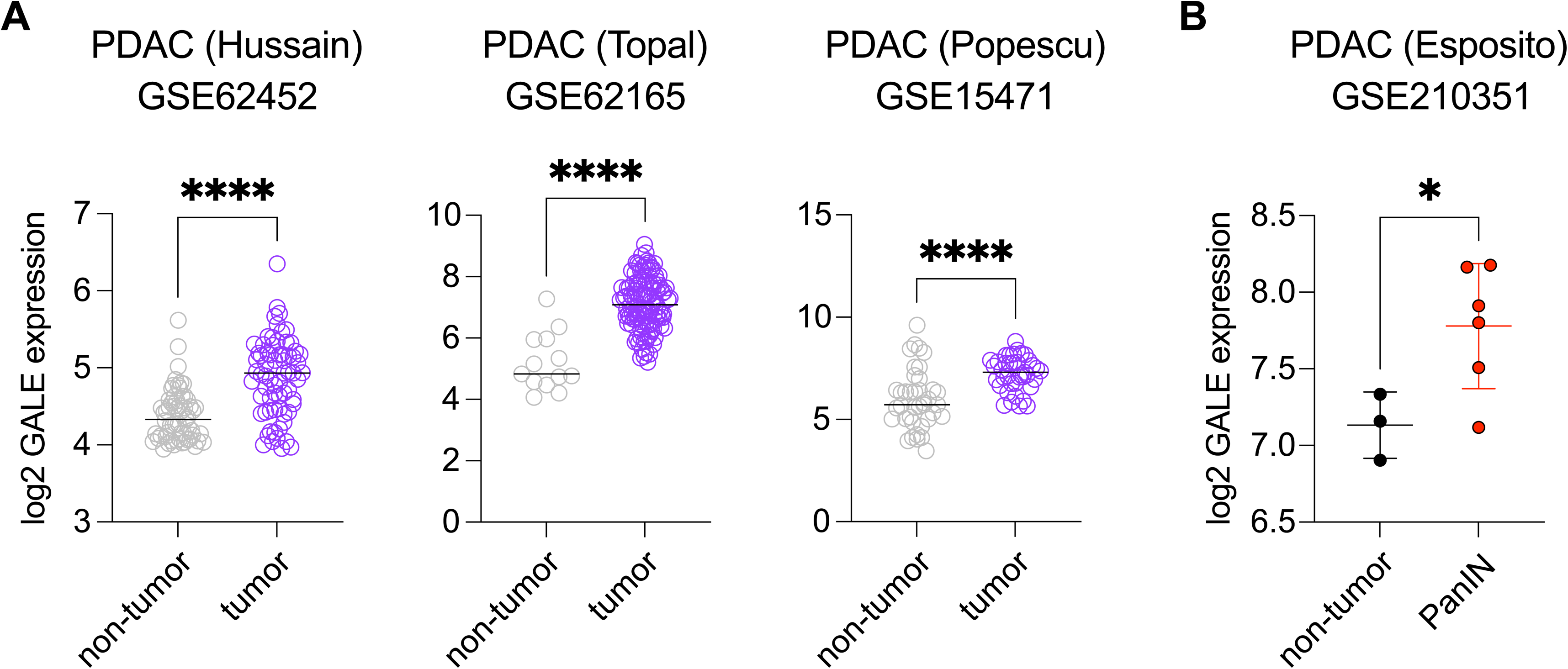
(A) *GALE* expression in tumor samples of PDAC in comparison to non-tumor tissue, obtained from GSE62452 (Hussain cohort), GSE62165 (Topal cohort), and GSE15471 (Popescu cohort), accessed via the R2 platform^26–28^. (B) *GALE* expression in pancreatic intraepithelial neoplasia (PanIN) lesions from PDAC patients, in comparison to non-tumor tissue, obtained from GSE210251 (Esposito cohort), accessed via the NCBI Gene Expression Omnibus (GEO) platform^29^. Unpaired, two-tailed *t test*; \**P* < 0.05, \*\**P* < 0.01, \*\*\**P* < 0.001, \*\*\*\**P* < 0.0001.

