## Supplementary material for "GALE-dependent glycoproteome remodelling is a determinant of oncogenic RAS transformation"

- Supplementary materials and methods
- Supplementary Table S1, S2, S3, S4, S5

**Supplementary Materials and Methods**

**Lentiviral transduction**

To generate lentiviruses, HEK293T cells were transfected with Calfectin (SignaGen, #SL100478) transfection reagent as well as a construct of interest and the psPAX2 (Addgene #12260), and pMD2.G (Addgene #12259) constructs, according to the manufacturer’s protocol. 48 hours after transfection, the viral supernatant was collected and filtered through a 0.45μm sterile PES filter unit (VWR, #76478-984). The lentivirus was stored at -80°C until use. CFPAC1 and H460 cells with stable repression of KRAS (shKRAS) were generated as previously reported^1^. PaTu-8988T shMYC cells were generated by infecting cells with relevant viral supernatant containing polybrene, for two rounds of 24-hour incubations, following which the cells were passaged and selected with 2 µg/mL puromycin for 7 days. Puromycin-resistant polyclonal cell populations were validated with qRT-PCR. NIH 3T3 KRASG12V and PANC-1 cells with stable repression of Gale/GALE (sgGale/GALE) were established by infecting cells with relevant viral supernatant containing polybrene, for two rounds of 24-hour incubations, following which the cells were passaged and selected with 5 µg/mL blasticidin for 7 days. Blasticidin-resistant polyclonal cell populations were validated with qRT-PCR.

**shRNA and CRISPR interference plasmids**

The shRNA construct targeting MYC were a kind gift from Celio Pouponnot (Institut Curie, France):CCGGACTGAAAGATTTAGCCATAATCTCGAGATTATGGCTAAATCTTTCAGTTTTTTG. CRISPRi constructs targeting mouse *Gale* were established by cloning sgRNA sequences corresponding to sgGale-1: GCATTCTACCGGCCGAACCTG and sgGale-2: GAGGGCTTAGTGAGCTCGGAA, or human *GALE* by cloning sgRNA sequences corresponding to sgGALE-1: GGTGCCTCTGCAGCAAGCGT and sgGALE-2: GTGCCTCTGCAGCAAGCGTG into Lenti-(BB)-EF1a-KRAB-dCas9-P2A-BlastR (Addgene plasmid #118154) while non-targeting sgEGFP construct corresponds to Lenti-(BB)-EF1a-KRAB-dCas9-P2A-BlastR EGFP-guide1 (Addgene plasmid #118158), both gifts from Jorge Ferrer (Centre for Genomic Regulation, Spain).

**siRNA transfection**

For siRNA knockdown experiments, cells were transfected at the time of seeding into six-well plates with individual or pooled siRNA or non-targeting scrambled siRNA at a final concentration of 25 nM using SilentFECT (BioRad, #1703362) according to the manufacturer’s instructions. Non-targeting scrambled siRNA (Dharmacon): UAAGGCUAUGAAGAGAUAC; Mouse Myc siRNA SMARTpool(Dharmacon): GACGAGACCUUCAUCAAGA, GACAGCAGCUCGCCCAAAU, GAAUUUCUAUCACCAGCAA, GUACAGCCCUAUUUCAUCU; Human MYC individual siRNA (Dharmacon), AACGUUAGCUUCACCAACA or CUACCAGGCUGCGCGCAAA.

**Quantitative RT-PCR**

Total RNA was extracted using the Direct-zol RNA Miniprep kit (Zymo Research, #R2052) according to manufacturer’s instruction. cDNA was reverse transcribed from RNA using High-Capacity cDNA Reverse Transcription Kit (Applied Biosystems, #4368813) according to manufacturer’s instruction. The cDNAs were quantified by real-time PCR analysis using SYBR Green Master Mix (Applied Biosystems, #4364344). The primer sequences are as follows:

| Target Gene | Forward (5’-3’) | Reverse (5’-3’) |
| --- | --- | --- |
| *Rplp0* | TAAAGACTGGAGACAAGGTG | GTGTACTCAGTCTCCACAGA |
| *GusB* | AAAATCACCCTGCGGTTGT | TGTGGGTGATCAGCGTCTT |
| *Gale* | CCATAACGCCATTCGTGGAG | TCCAGAGGCTTCTGCACTGA |
| *Myc* | ATGCCCCTCAACGTGAACTTC | CGCAACATAGGATGGAGAGCA |
| *Cdc20* | TTCGTGTTCGAGAGCGATTTG | ACCTTGGAACTAGATTTGCCAG |
| *Aurka* | CTGGATGCTGCAAACGGATAG | CGAAGGGAACAGTGGTCTTAACA |
| *RPLP0* | GGCGACCTGGAAGTCCAACT | CCATCAGCACCACAGCCTTC |
| *GUSB* | GTTTTTGATCCAGACCCAGATG | GCCCATTATTCAGAGCGAGTA |
| *GALE* | CTGGAGGCTGGCTACTTGC | CCCTGGTCCAAAATGTCCATCT |
| *MYC* | GTCAAGAGGCGAACACACAAC | TTGGACGGACAGGATGTATGC |
| *CDC20* | GACCACTCCTAGCAAACCTGG | GGGCGTCTGGCTGTTTTCA |
| *KRAS* | CAGTACAGTGCAATGAGGGAC | CCTGAGCCTGTTTTGTGTCTAC |

**Reagents**

Blasticidin (Invivogen GmbH, #ant-bl-05), AMG510 (MedChemExpress, #HY-114277), PD184352 (Cayman, #11580), MK2206 (Cayman, #11593), 10058-F4 (MedChemExpress, #HY-12702), N-Acetyl-D-galactosamine (GalNAc) (Biozol, #CBS-MA04390), galactose (Sigma, #G0750).

**Soft agar colony formation assays**

Cells were plated in 6-well plates with 8,000 cells per well in DMEM 10% FBS or DMEM 10% bovine calf serum in a top layer of 0.25% agar added over a base layer of 0.4% agar in DMEM 10% FBS or DMEM 10% bovine calf serum. Where indicated, supplementation with galactose or GalNAc were added to the top agar layer. Cells were fed once every three days with 1 ml of corresponding medium onto the top layer, and containing relevant sugars or vehicle if necessary. After 2-3 weeks at 37°C, colonies were stained with MTT following which they were imaged and quantified using ImageJ software. The percentage of colony forming cells was calculated.

**Chorioallantoic membrane (CAM) model**

For CAM xenografts, specific-pathogen-free chicken (SPF) eggs freshly fertilized (d0) and supplied by VALO BioMedia GmbH, Germany, were used. Cells were washed with PBS, trypsinized, centrifuged and resuspended in PBS at a concentration of 0.5-1*10^6^/50µL PBS per cell line. 50 µL PBS was inoculated onto the CAM at day 10 within a sterile silicon ring. Eggs were incubated at 37°C for 7 days in a stationary incubator. Where indicated, cells were incubated in appropriate cell culture media containing galactose or GalNAc, and three days after inoculation of the cells, tumors were treated by pipetting 50µl of galactose or GalNAc-containing media onto the CAM within the sterile silicon ring. Tumors were harvested and weighed 7 days following inoculation, embryos were immediately sacrificed. All experimental procedures involving avian embryos were conducted in strict accordance with institutional guidelines and established standard operating procedures (SOPs) for the care and use of laboratory animals**.**

**Mouse tumorigenicity assays**

For NIH 3T3 KRAS^G12V^ sgGale and sgEGFP cells, aliquots of 0.5 × 10^6^ cells were resuspended in 100 μL of PBS and injected subcutaneously into the flanks of 5- to 6-week-old female NOD *Scid* Gamma mice using standard procedures. When tumours became palpable, mice were evaluated for tumor growth every 2 days until the experimental humane endpoints. Tumors were measured with a caliper, and volumes were estimated using the following formula: tumor length × (tumor width)2 × π/6 mm3. All animal experiments underwent ethical approval from the Animal Care Committee of the University of British Columbia (License number: A21-0209).

For PANC1 sgGALE and sgEGFP cells, suspensions of 0.5 × 10^6^ in 100 µL of PBS: Matrigel 1:1 (VWR, #734-1101) were implanted subcutaneously in the flank of athymic Foxn1nu/nu mice (female, 7 weeks, Janvier labs, 8 mice per group). Tumors were allowed to reach 50 mm^3^ before serial monitorization of tumor growth and body weight (2-3 times per week). Tumor dimensions were measured using a digital Vernier caliper, and volume was calculated as follows: volume = (width^2^ × length)/2. Mice were euthanized when tumors reached maximal length of 14 mm or volume reached 1000 mm³, to comply with maximal tumor size allowed by the ethics committee. Tumors were then removed and sectioned, and pieces were fixed in 10% formalin or frozen for analysis. Animal procedures were conducted at Aarhus University´s animal facility in accordance with institu-tional and national regulations and approved by the Danish Animal Experiments Inspectorate (License number: 2024-15-0201-01741). Animals were housed under controlled temperature and hu-midity with ad libitum access to food and water and a 12 h light–dark cycle. All mice were female and sex was not a specified variable for analysis in this study.

**Western blotting**

Cells were lysed in RIPA buffer (150 mM NaCl, 50 mM Tris-HCl, pH 8, 1% Triton X-100, 0.5% sodium deoxycholate, and 0.1% SDS) supplemented with cOmpleteTM, EDTA-free Protease Inhibitor Cocktail (Sigma, #11873580001) and phosphatase inhibitors (PhosphoSTOP, Roche, #4906845001). Cell lysates were incubated for 30 minutes on ice, centrifuged at 14,000 x g for 15 min at 4°C following which supernatant fractions were collected. Protein concentration and normalization was performed using PierceTM BCA Protein Assay Kit (Thermo Fisher Scientific, #23225) according to manufacturer’s protocol. Protein lysates were resolved by SDS-PAGE and transferred to nitrocellulose membranes (GE Healthcare), 10402096. Membranes were blocked with 5% BSA TBS-Tween (20 mM Tris-HCl, pH 7.4, 150 mM NaCl, 0.1% Tween 20) and probed with the primary antibodies listed in Supplementary table 2. Secondary anti-mouse (Li-Cor, #926-32210) or anti-rabbit (Li-Cor, #926-32211) antibodies were used and fluorescent signal was detected with the LI-COR Odyssey CLx system. The following antibodies were used:

| Antibody | Dilution | Manufacturer |
| --- | --- | --- |
| AKT (pan, C67E7) | 1:1000 | Cell Signaling Technology, #4691 |
| c-MYC (D84C12) | 1:1000 | Cell Signaling Technology, #5605 |
| ERK (p44/42 MAPK (Erk1/2), (137F5) | 1:1000 | Cell Signaling Technology, #4695 |
| GALE (OTI1C4) | 1:1000 | Abcam, #118033 |
| GAPDH | 1:1000 | Cell Signaling Technology, #2118 |
| p-AKT (Ser473) (D9E) XP | 1:1000 | Cell Signaling Technology, #4060 |
| p-ERK (Thr202/Tyr204) | 1:1000 | Cell Signaling  Technology, #9101 |
| S6 ribosomal protein (5G10) | 1:1000 | Cell Signaling Technology, #2217 |
| Vinculin | 1:1000 | Cell Signaling Technology, #4650 |

**Immunohistochemistry of tissue microarray**

Tissue microarrays were constructed from 240 PDAC resection specimens obtained between 1995 and 2017. Clinicopathological data were retrieved from institutional records and updated in accordance with the most current guidelines. For each case, representative areas were selected from the corresponding resection specimen and arrayed as tissue cores measuring 1 mm in diameter. When available, each case was represented by two cores from the central tumor area, one core from the peripheral tumor area, two cores from lymph node metastases, and one core from non-neoplastic pancreatic tissue.

Immunohistochemical staining for GALE (Abcam, #118033) was performed manually using human gastric tissue as a positive control. Tissue sections were deparaffinized overnight at 60 °C, treated with xylene for 20 min, and rehydrated through graded ethanol to distilled water. Antigen retrieval was performed in citrate buffer, pH 6, using a pressure cooker for 15 min. Endogenous peroxidase activity was blocked with 3% H₂O₂ for 10 min.

Sections were circumscribed with a PAP pen and incubated with the primary anti-GALE antibody diluted 1:5000 in blocking solution containing bovine serum albumin, goat serum, and Triton X for 1 h in a dark humidified chamber. After washing with TBS-T, sections were incubated with anti-mouse IgG secondary antibody from the VECTASTAIN® ABC-HRP Kit for 30 min, followed by incubation with the avidin–biotin complex for 30 min. Immunoreactivity was visualized with diaminobenzidine (DAB) for 3 min. Sections were counterstained with Mayer’s hematoxylin, dehydrated through graded ethanol, cleared in xylene, and mounted with Eukitt®.

***C. elegans* multivulva phenotype experiment**

RNA was isolated from adult stage N2 *Caenorhabditis elegans* (RNeasy kit, QIAGEN, #74106). *gale-1* cDNA was amplified from total RNA using the Superscript IV reverse transcriptase (Thermo) and cloned with restriction digest and ligation into the L4440 RNAi plasmid. Following sequence verification, competent HT115 *E. coli* were transformed with the L4440:: *gale-1* plasmid for use in RNAi experiments.

HT115 E. coli expressing *gale-1* double-stranded RNA were grown overnight at 37 °C in LB medium supplemented with ampicillin (100 μg/mL) and tetracycline (10 μg/mL). Cultures were induced with 0.4 mM IPTG for 4 h, concentrated 10-fold by centrifugation and seeded on nematode growth media (NGM) plates containing 0.25 mM IPTG and 25 μg/mL carbenicillin. HT115 *E. coli* carrying the empty L4440 vector were used as controls for RNAi experiments.

The *C.* *elegans* strain MT2124, *let-60(n1046)*, was used in this study. The n1046 allele is a gain-of-function mutation in the *C. elegans* Ras homolog *let-60*, containing a glycine-to-glutamic acid substitution at amino acid 13 (G13E). Synchronized embryos obtained by the bleaching method. Briefly, a plate of gravid hermaphrodite worms grown to near starvation on 60 mm plates were collected in 1.5 ml Eppendorf tubes with M9 buffer + 0.1%Triton X-100 and pelleted for 1 minute at 500 rpm. Buffer was carefully aspirated from the pelleted worms and replaced with bleach solution (825 μl of Milli-Q Water + 375 μl of 1M NaOH + 300 μl household bleach) with gentle agitation. Embryos were pelleted by centrifugation at 5,000 rpm for 30 s and bleach solution aspirated and replaced with 1 ml of M9 buffer, then pelleted at 5,000 rpm. Buffer was aspirated from the pelleted eggs and washed two more times with M9. Final pelleted eggs were resuspended in 500 l of M9 and incubated on a rotator overnight until they hatched and arrested at the first larval stage (L1). The next morning L1 stage worms were pipetted on RNAi bacterial lawns (50–100 worms per condition) and maintained at 20 °C for 2–3 days until the developed to adulthood. Animals were scored for the multivulva (Muv) phenotype as previously described (Subramanian et al., 2021). Worms displaying at least one pseudovulva were classified as Muv, and percent Muv was calculated as the number of Muv animals divided by the total number of adult animals scored per plate.

### Glycoproteome analysis via SAX-ERLIC enrichment and mass spectrometry

Glycopeptides were enriched from cell pellets sequentially by strong-anion-exchange followed by electrostatic-repulsion-hydrophilic interaction chromatography (SAX-ERLIC), as previously described^2,3^. Enriched glycopeptides were analyes on an ECLIPSE Trybrid system (ThermoFischer) using combined stHCD-pd-EthcD fragmentation in data dependent acquisition mode, method adapted from Sutherland et al., 2024^4^. Data analysis was performed using Fragpipe (v23.0). Glycoproteome data was searched against either a human protein database containing 20453 protein sequences (Uniprot, UP000005640, downloaded on 13/01/2026) or a mouse database containing 17352 protein sequences (Uniprot, UP000000589, downloaded on 03/11/2025). The N-glycan search was performed using the built-in workflow with the following changes: MSFragger: allowed up to 3 missed cleavages, and a glycan database containing 1709 glycans with allowed glycan modifications (up to 2 x NH4+, 1 x Na) bringing the total number of glycans to 6830, the database was extracted from Riley et al., 2019^5^. For O-glycan analysis, the built-in workflow “glyco-O-Hybrid” search was used, using the same modifications as above, but using the also built.in “Oglyc78” glycan library, with the same glycan modes as above. The data was then post-processed using in-house R Scripts.

**Bioinformatics analyses of gene expression patterns and Kaplan Meier survival analyses in human tissue samples**

For gene expression analysis, RNA-seq data from TCGA and the GTEx projects were analyzed with UCSC Xena platform.

Kaplan Meier analyses were performed using KM-Plotter software (https://kmplot.com/analysis/) using default parameters, from either gene-chip data (colon adenocarcinoma, gastric adenocarcinoma, liver hepatocellular carcinoma, glioma, multiple myeloma, ovarian carcinoma, and sarcoma) or RNA sequencing data (pancreatic adenocarcinoma and rectal adenocarcinoma)^6^. For each tumor type, GALE expression was stratified by median *GALE* expression except in live hepatocellular carcinoma, where the cut-off was first-quartile. All analyses were performed on 24/01/2026.

**Single cell RNA sequencing data analysis**

Single cell RNA-seq data for pancreatic ductal adenocarcinoma (PDAC) patients and PDAC mouse model were collected from previously published research papers^7,8^. The datasets for human and mice are available in the Genome Sequence Archive (GSA) under the accession number: CRA001160 for Peng et al., 2019 and in the Gene Expression Omnibus (GEO) repository under the accession number: GSE141017 for Schlesinger et al., 2020. The analysis was orchestrated in R v4.4.3. The gene-barcode matrices with gene expression counts (rows) and barcoded cells (columns) were imported into the Seurat R package (v5.3.0). All functions were run with default parameters, if not stated otherwise. In both datasets, cells were retained based on quality thresholds: 200 to 6000 expressed genes and mitochondrial gene expression (< 15% for Peng et al., 2019 and < 5% for Schlesinger et al., 2020). Additionally, only genes expressed in at least three cells were included.

The preprocessed matrix was normalized with the *NormalizeData* function of Seurat, and the top 2000 variable features were selected with the variance stabilizing transformation (vst) of the *FindVariableFeatures* function. The gene expression was scaled with the *ScaleData* function and principal component analysis (PCA) was performed with *RunPCA*. Based on the elbow plot and Seurat’s PCHEatmap for the first 50 dimensions, the top principal components (*n*=10 PCs in Peng et al., 2019 and *n*=13 PCs in Schlesinger et al., 2020) were selected. Non-linear dimensionality data reduction was performed with Uniform Manifold Approximation and Projection (UMAP) and t-distributed Stochastic Neighbor Embedding (t-SNE), and the data was visualized in a two-dimensional space. The shared nearest neighbors (SNN) for both datasets were computed using the *FindNeighbors* function of Seurat with the default k parameter (k=20). To determine the optimal resolution for clustering, a variety of resolutions ranging from 0.1 to 1.2 in increments of 0.2 were systematically evaluated. For each resolution, an unsupervised graph-based *Leiden* clustering (algorithm=4) was run using Seurat’s *FindClusters* function. The optimal resolution was selected based on the iterative process of the *clustree* [9] analysis (*r*=0.7 in ^4^ and *r*=0.5 in ^3^), and the cells were clustered into specific groups (*n*=25 clusters in Peng et al., 2019 and *n*=20 clusters in Schlesinger et al., 2020).

A subset of PDAC-associated markers in mice (*Kras*, *Trp53*, *Smad4*, *Cdkn2a*, *Thy1*, *Id1*, *Krt19*, *Ceacam6*, *Muc1*, *Muc5ac*, *S100p*, *Msln*, *Tff1*, *Tspan8*, *Mmp7*, *Anxa10*, *Krt7*, *Epcam*, *Gale*, *Myc*, *Cpa1*, *tdTomato*, *Foxq1*, *Onecut2*, *Prox1*, *Sox9*, *Rbpjl*, *Id3*, *Runx1*, *Gkn1*, *Gkn2*, *Muc6*, *Pga5*, *Pgc*, *Dclk1*, *Mki67*, *Tgfbl*, *Ins1*) was used to check for their presence in all clusters. Markers of interest (*Kras*, *Myc*, *Gale*, and *tdTomato* and *KRAS*, *MYC*, *GALE*) were visualized on a separate layer on top of the UMAP projection. Violin plots and ridge plots were generated for each of the expressed markers. Plots for additional present markers in the respective datasets can be found in the Appendix. In the human dataset, the respective orthologous mouse genes retrieved with the *convert_mouse_to_human_symbols* function from the *nichenetr* R package [10] have been used to validate their presence across all clusters.

To classify potential malignant clusters, a module score was assigned for each cell based on the list of present PDAC markers using the *AddModuleScore* function of Seurat and a cluster-wise mean PDAC score was computed. The clusters were sorted in decreasing order by the mean PDAC score and the top *n* clusters (*n*=4 clusters in Peng et al., 2019 and *n*=2 clusters in Schlesinger et al., 2020) were classified as potentially malignant. These clusters were visualized along with the reference cells in a violin plot and a UMAP with the *VlnPlot* and *DimPlot* functions of Seurat respectively. Differentially expressed genes were identified by applying the *FindAllMarkers* function of Seurat. The one-vs.-rest Wilcoxon signed-rank statistical test (two-sided non-parametric test) was run with the parameters *logfc.threshold=0.25* and *min.pct = 0.25*. The top *n*=50 up and down regulated expressed markers for each malignant cluster were identified by filtering based on a p-adjusted value < 0.05, and sorted by the log2 fold change in decreasing order. In human and mice, the cells were annotated with reference markers (Ductal cell 1: *MBP*, *CFTR*, *MMP7*; Ductal cell 2: *KRT19*, *KRT7*, *TSPAN8*, *SLPI*; Acinar cells: *PRSS1*, *CTRB1*, *CTRB2*, *REG1B*; Endocrine cells: *CHGB*, *CHGA*, *INS*, *IAPP*; Stellate cells: *RGS5*, *ACTA2*, *PDGFRB*, *ADIRF*; Fibroblasts: *LUM*, *DCN*, *COL1A1*; Endothelial cells: *CDH5*, *PLVAP*, *VWF*, *CLDN5*; Macrophages: *AIF1*, *CD64*, *CD14*, *CD68*; T cells: *CD3D*, *CD3E*, *CD4*, *CD8*; and B cells: *MS4A1*, *CD79A*, *CD79B*, and *CD52* in Peng et al.; Neuroendocrine cells: *Resp18*, Pericytes: *Rgs5*, Immune cells: *Ptprc*, Acinar cells: *Cpa1*, Ductal, metaplastic, and tumor cells: *Krt19*; Fibroblasts: *Col1a1*; and Endothelial cells: *Pecam1*).

**Quantification and Statistical Analysis**

All experiments were, if not otherwise stated, independently carried out at least three times. Statistical significance was calculated using Student’s t-test in GraphPad Prism 8. The data are represented as means +/- standard deviation. A p-value of less than 0.05 was considered to be significant.
